# A eukaryote-specific factor mediates an early step in the assembly of plant photosystem II

**DOI:** 10.1101/2023.05.12.540541

**Authors:** Jakob-Maximilian Keller, Maureen Julia Frieboes, Ludwig Jödecke, Sandrine Kappel, Natalia Wulff, Tobias Rindfleisch, Omar Sandoval-Ibanez, Ines Gerlach, Wolfram Thiele, Ralph Bock, Jürgen Eirich, Iris Finkemeier, Danja Schünemann, Reimo Zoschke, Mark Aurel Schöttler, Ute Armbruster

**Author notes:** **Correspondence:** Ute Armbruster, Institute of Molecular Photosynthesis, Heinrich Heine University Düsseldorf, Universitätsstraße 1, 40225 Düsseldorf, Germany.

## Abstract

The initial step of oxygenic photosynthesis is the thermodynamically challenging extraction of electrons from water and the release of molecular oxygen. This light-driven process, which is the basis of life on Earth, is catalyzed by the photosystem II (PSII) within the thylakoid membrane of photosynthetic organisms. The biogenesis of PSII requires a controlled step-wise assembly process of which the early steps are considered to be highly conserved between plants and their cyanobacterial progenitors. This assembly process involves auxiliary proteins, which are likewise conserved. In the present work, we show that in plants, the early assembly step, in which the PSII reaction center (RC) is associated with the intrinsic antenna protein CP47 to form the RC47 intermediate, is facilitated by a novel eukaryote-exclusive assembly factor. This factor, we named DEAP2 for DECREASED ELECTRON TRANSPORT AT PSII, works in concert with the conserved PAM68 assembly factor. The *deap2* and *pam68* mutants showed similar defects in PSII accumulation and assembly of the RC47 intermediate. The combined lack of both proteins results in a loss of functional PSII and the inability of plants to grow photoautotrophically on soil. While overexpression of DEAP2 partially rescued the *pam68* PSII accumulation phenotype, this effect was not reciprocal. DEAP2 accumulates at 20-fold higher levels than PAM68, together suggesting that both proteins have distinct functions. In summary, our results uncover eukaryotic adjustments to the PSII assembly process, which involve the addition of DEAP2 for the rapid progression from RC to RC47.

## Introduction

Life on earth is shaped by the continuous production of molecular oxygen by photosynthetic organisms. Molecular oxygen is released during the initial step of photosynthesis, in which light energy is used to extract electrons from water. This thermodynamically demanding reaction is catalyzed by the photosystem II (PSII) multi protein complex. Under natural light conditions, plant PSII undergoes continuous damage, disassembly and repair (Komenda et al., 2012; Nickelsen and Rengstl, 2013; Lu, 2016). A mechanistic understanding of PSII biogenesis and repair in plants is pivotal on multiple levels. Besides providing fundamental insight into plant’s bioenergetic blueprint, it can stipulate strategies to enhance crop yield by optimizing photosynthesis or support synthetic biology-based methods to harness light energy.

PSII constitutes a light-driven oxidoreductase localized in the photosynthetic thylakoid membrane, which oxidizes water at the lumen side and via a sequence of redox reactions reduces the membrane integral plastoquinone carrier at the stromal side. This life-defining molecular machine contains nuclear and plastid-encoded protein subunits and in the functional form of a plant supercomplex consists in total of more than 500 components (Shen, 2015; Wei et al., 2016). All these constituents have to be assembled in a concerted fashion to obtain a functional complex. Multiple co-factors, which either absorb light energy such as chlorophylls or are highly reactive such as hemes and the manganese cluster strictly require integration simultaneously with the protein assembly process. Thus, a highly organized and stepwise assembly procedure is fundamental to avoid early damage of the nascent protein complex by the formation of highly reactive intermediates (Zagari et al., 2017). This involves the function of molecular chaperones, which are referred to as PSII assembly factors.

In the early steps of PSII biogenesis, the core proteins are assembled to form the reaction center (RC) assembly intermediate. This intermediate consists of the cytochrome *b_559_* forming PsbE and F, the low-molecular-mass PsbI and the D1 and D2 subunits, which bind most redox-active cofactors of rapid electron transport within PSII (reviewed (Komenda et al., 2012; Nickelsen and Rengstl, 2013; Lu, 2016). Then, the addition of the intrinsic antenna CP47 and PsbH, PsbR, and PsbTc leads to the formation of the so called RC47 assembly intermediate (Komenda et al., 2004; Rokka et al., 2005). Subsequently, the intrinsic antenna CP43, multiple small membrane-intrinsic subunits, and the extrinsic subunits PsbO, PsbP, and PsbQ, which stabilize the oxygen evolving complex, are added to derive the PSII monomer. Ultimately, two PSII monomers form a PSII dimer and bind additional light harvesting complexes to give rise to PSII supercomplexes (SC, (Shevela et al., 2016)).

The first assembly steps up to RC47 proceed in plant chloroplast in a very similar way as in cyanobacteria. All known assembly factors involved in the formation of RC47 in plants have orthologs in cyanobacteria, further arguing for the evolutionary conservation of these early steps of the assembly process (Nickelsen and Rengstl, 2013; Lu, 2016). In *Arabidopsis thaliana* (Arabidopsis), RC assembly requires multiple essential factors that facilitate thylakoid membrane targeting, protein insertion and folding, as well as assembly. Mutants that lack these factors cannot form functional PSII and thus are unable to grow photoautotrophically (Meurer et al., 1998; Link et al., 2012; Lu, 2016; Myouga et al., 2018; Li et al., 2019). Plants that lack LOW ACCUMULATION OF PSII (LPA1) can still assemble some functional PSII, but levels are strongly decreased, which leads to severely delayed growth compared with wild type (WT) on soil (Peng et al., 2006). LPA1 was shown to associate with nascent D1 to promote the formation of the RC intermediate (Peng et al., 2006; Armbruster et al., 2010). For the next step in PSII assembly, which is the progression from RC to RC47, so far only one specific assembly factor has been described, which is the PHOTOSYNTHESIS AFFECTED MUTANT (PAM68) protein (Armbruster et al., 2010). Plants lacking PAM68 display a slowing down of PSII assembly directly after the RC intermediate. This leads to a similar phenotype as in the *lpa1* mutant with low PSII accumulation and slow growth (Armbruster et al., 2010). The cyanobacterial ortholog of PAM68 from Synechocystis was found to interact with ribosomes and promote CP47 synthesis and it was suggested that PAM68 is involved in the co-translational insertion of chlorophyll into CP47 (Bucinska et al., 2018). In Arabidopsis however, no differences could be observed for the synthesis of CP47 between *pam68* mutants and WT (Armbruster et al., 2010), questioning whether this function of PAM68 is conserved between plants and cyanobacteria.

In the present study, we discovered the first plant assembly factor for the early stages of PSII biogenesis not conserved in cyanobacteria and named it DEAP2 (DECREASED ELECTRON TRANSPORT AT PSII). We demonstrate that DEAP2 acts in concert with the evolutionary conserved PAM68 to facilitate the progression of PSII assembly from the RC to the RC47 intermediate.

## Results

### Lack of DEAP2 affects plant growth, pigmentation and PSII function

During an initial screen of *Arabidopsis thaliana* (Arabidopsis) T-DNA insertion lines under fluctuating light conditions, we found that the mutants of the line SALK_048033 grew very slowly and had chlorotic leaves (Supplemental Fig. S1). When grown under low non-fluctuating light conditions, these mutants still showed a strong growth retardation and paler leaves. Salk_048033 carries a T-DNA insertion in the gene *At3g51510* (Fig. 1A). To confirm that *At3g51510* was the casual gene for the observed phenotype, two more mutants were generated via CRISPR-Cas gene editing. Under low light conditions, all plants with lesions in *At3g51510* showed the same slow growth and pale leaf phenotype, which was complemented by the introduction of the *At3g51510* cDNA expressed from the 35S promoter into SALK_048033 (Fig. 1B, Supplemental Fig. S2A). All three independent *at3g51510* mutants showed a strong decrease in the maximum quantum efficiency of PSII in the dark-adapted state (Fv/Fm) and were thus termed *decreased electron flow at PSII* (*deap2*) with the two analyzed complemented lines being referred to as *cDEAP2-1* and *-2* (Fig. 1B). We confirmed the absence of DEAP2 protein in all three *deap2* lines by generating a specific antibody against the 18 most C-terminal amino acids of the protein (Supplemental Fig. S2B, C). While no protein was detected in the mutants, an upward shift of the signal was detected in the *cDEAP2* lines compared with the WT. The *cDEAP2* lines carry a Myc tag at the C-terminal end of DEAP2, which adds 1.2 kDa to the protein. The growth, chlorophyll (Chl) content, and Fv/Fm phenotypes of *deap2* mutants were similar to that of the previously reported *pam68* mutant, which is defective in the assembly of the RC47 intermediate of PSII ((Armbruster et al., 2010), Fig. 1C-F). The reduced Chl a/b ratio indicated a stronger loss of photosystem reaction centers (binding only Chl a) relative to antenna proteins (binding both Chl a and b) in the two mutants. By analyzing the PSI donor side limitation under growth light intensity, we found that it was severely increased to the same degree in *deap2* and *pam68* compared with WT and *cDEAP2*. This result showed that linear electron transport towards PSI was decreased, supporting that activity of PSII was more negatively affected than PSI in both mutants (Fig. 1G).

**Figure 1:**
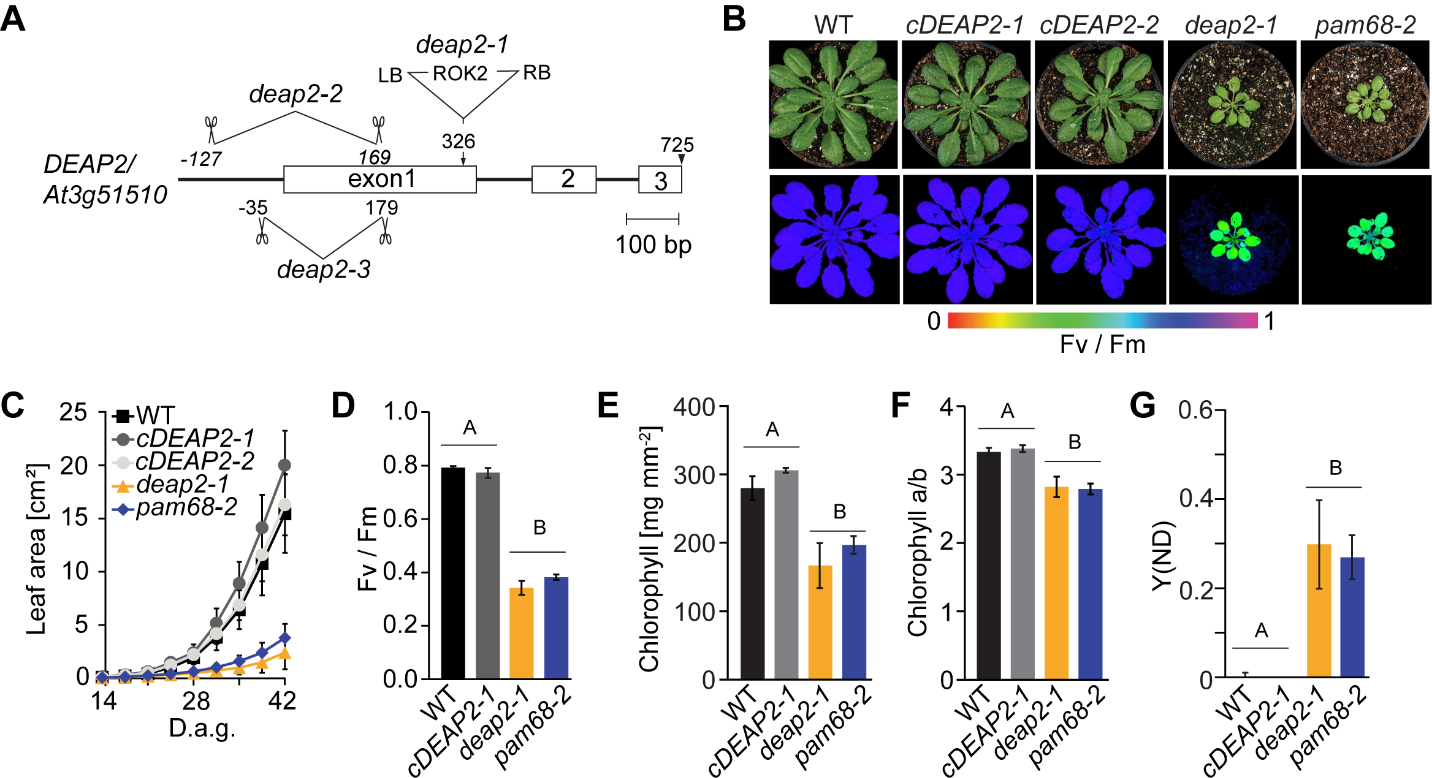
The *deap2* mutant shows a defect in growth, pigmentation and PSII function comparable to the PSII-assembly factor deficient *pam68*. **(A)** Map of the gene locus *DEAP2/At3g51510*. Exons are displayed as white boxes, introns as black lines. Location of the T-DNA insertion in *deap2-1* (SALK_048033) and the deletions obtained by CRISPR/Cas, *deap2-2* and *deap2-3* are shown. Deletions are indicated by scissors giving the cutting site relative to the start codon as determined by sequencing of genomic DNA from these lines. **(B)** Pictures (top panel) and PSII maximum quantum yield (Fv/Fm) false color images taken with the Imaging-PAM (bottom panel) of eight-week old Columbia-0 (wild type, WT), two independent complementation lines, *cDEAP2-1* and *cDEAP2-2*, *deap2-1* and *pam68-2* (top panel) grown at 50 µmol photons m^−2^ s^−1^ with a 12/12 h light/dark cycle. Signal intensities for Fv/Fm are given by the false color scale below the panels. **(C)** Increase in leaf area of the different genotypes measured from 14 to 42 days after germination (d.a.g.). Shown is the average of n=10 ±SD. **(D-G)** Bar graphs of (D), chlorophyll content per leaf area (E), Chlorophyll a/b ratio (F) and yield of the PSI donor site limitation at growth light intensity (G). Average is shown of n=5-9 ±SD. The capital letters above the graphs denote the two significantly different groups (p < 0.05) as determined by one-way ANOVA and Holm-Sidak multiple comparison test.

### DEAP2 is a eukaryote-specific thylakoid membrane protein found only in the green lineage

For localization, the C-terminus of DEAP2 was fused to YFP and the respective construct was transiently expressed in leaves of *Nicotiana benthamiana*. This experiment revealed that DEAP2 colocalizes with Chl a fluorescence in the chloroplast (Fig. 2A). To further investigate the localization of DEAP2, we isolated intact chloroplasts from Arabidopsis WT rosettes and fractionated them into envelope and thylakoid membranes as well as soluble stroma. Immunoblot analysis using the specific DEAP2 antibody confirmed the exclusive localization of DEAP2 in the thylakoid membrane together with the Chl-binding light harvesting protein Lhcb1 (Fig. 2B, lower panel). Treatment of WT thylakoids with chaotropic salts revealed that only little of the DEAP2 protein could be extracted, which is consistent with this protein constituting a thylakoid integral protein (Fig. 2C), thus supporting the prediction of two transmembrane helices (Fig 2D).

**Figure 2:**
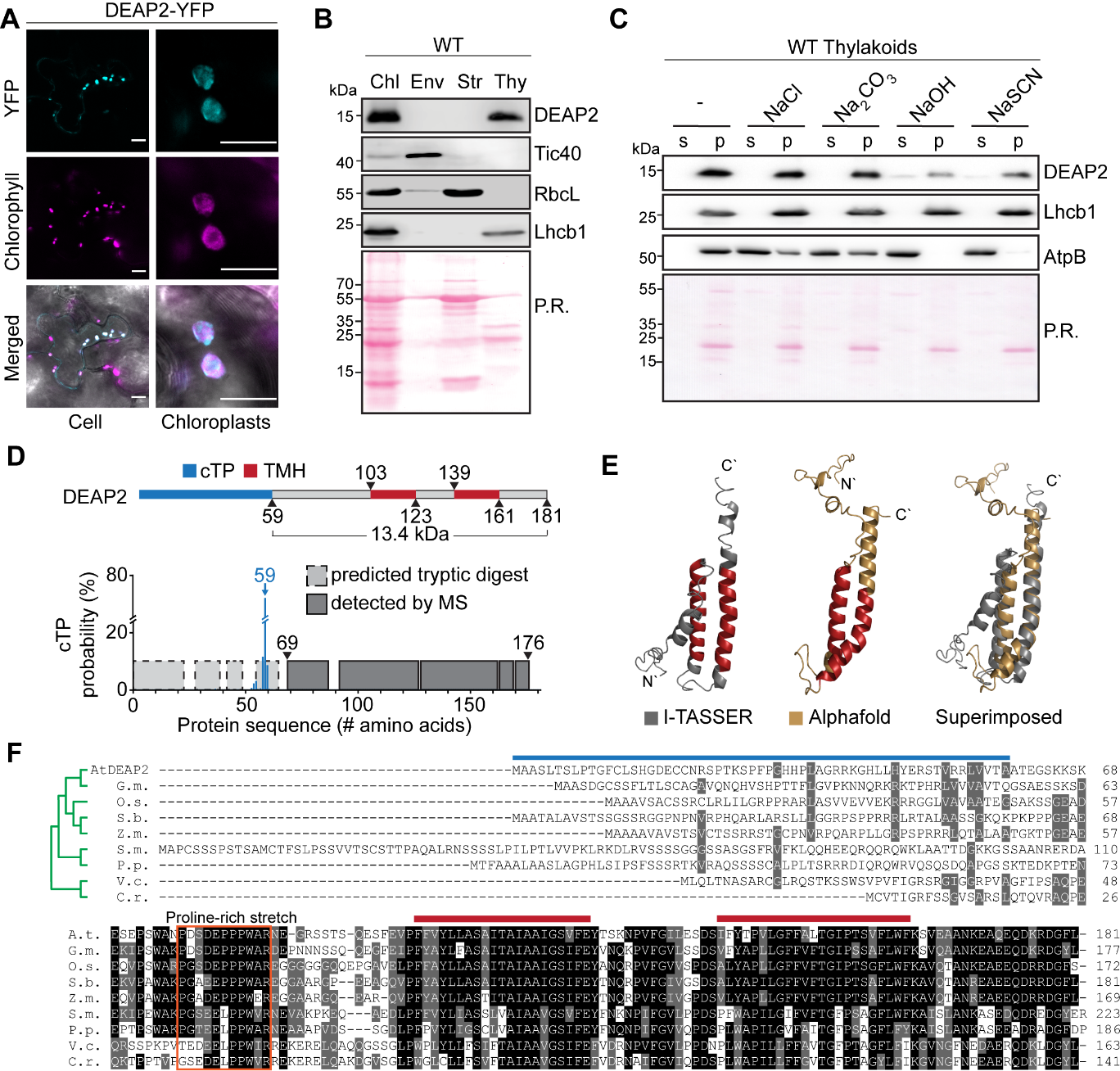
DEAP2 is a thylakoid integral protein conserved in the eukaryotic green lineage. **(A)** Localization of YFP-tagged DEAP2 in *Nicotiana benthamiana* by fluorescence microscopy. The individual signals of chlorophyll in magenta (top) and YFP in cyan (middle) and a merged image (bottom) of a cell (left) and individual chloroplasts (right) are shown. Scale bar represents 25 µM and 10 µM on left and right, respectively. **(B)** Immunoblots using the specific DEAP2 antibody show the protein localizes in the thylakoid membrane: proteins of whole chloroplasts (Chl), envelope (Env), stroma (Str) and thylakoids (Thy) were probed for DEAP2, Tic40 as a control for the envelope membrane, RbcL as a control for the stroma and Lhcb1 as a control for the thylakoid membrane. **(C)** WT thylakoids were incubated either with buffer only (-) or added NaCl, Na2CO3, NaOH or NaSCN and subsequently separated into pellet (p) and supernatant (s) fractions by centrifugation. Immuno-detection was carried out on fraction using antibodies against DEAP2, Lhcb1 as a membrane integral control and AtpB as a membrane associated control. (B-C) Ponceau staining (P.R.) of the membrane prior to immunodetection is shown. Numbers left of the immunoblots indicate molecular weight in kDa of protein standards. **(D)** Protein model of DEAP2 with predicted chloroplast targeting peptide (cTP) and transmembrane helices (TMH). cTP cleavage site prediction by TargetP was supported by lack of MS-peptides in mature DEAP2 enriched from the thylakoid membrane of *cDEAP2-1* by using a Myc-trap. **(E**) Protein structure of mature DEAP2 predicted by AlphaFold and I-TASSER and both structures superimposed. **(F)** Sequence alignment and phylogenetic reconstructions (green tree) of DEAP2 homologs of *Glycine max* (G.m.), *Oryza sativa* (O.s.), *Sorghum bicolor* (S.b.), *Zea mays* (Z.m.), the mosses *Sphagnum maghellanicum* (S.m.) and *Physcomitrium patens* (P.p.), and the green algae *Volvox carteri* (V.c.) and *Chlamydomonas reinhardtii* (C.r.). 50% to <100% sequence similarity is marked grey and 100% black. Bars above sequences indicate cTP and TM as in (D). The orange box indicates a conserved proline-rich domain N-terminal of the transmembrane domains.

A predicted chloroplast transit peptide (cTP) cleavage site around alanine (A) 59 of the precursor protein was supported by MS analysis of enriched mature DEAP2 from the thylakoid membrane, which only detected peptides C-terminal of this site (Fig. 2C, Supplemental Table S1-S2). The molecular weight of the mature DEAP2 protein was predicted to be 13.4 kDa, consistent with the immunoblots, which showed that the WT and Myc-tagged DEAP2 proteins migrated just below or at the same height as the 15 kDa marker protein band, respectively (Fig. 2B). Protein structure predictions suggested both transmembrane helices (TMH) of the Arabidopsis DEAP2 to be connected via an unstructured region and to extend to the C-terminus as a helical structure (Fig. 2E). The DEAP2 protein is conserved in the green lineage of photosynthetic eukaryotes (Fig. 2F; (Karpowicz et al., 2011)). Besides a high conservation of the region encompassing the two TMHs and the C-terminus, the protein shows a conserved domain in the N-terminus of the mature protein, which is proline-rich with an increasing number of prolines during the evolution from algae to vascular plants.

### Loss of DEAP2 has negative consequences specifically for the accumulation of PSII

To obtain further insight into the accumulation of thylakoid protein complexes in the *deap2* mutant, we performed immunoblot analyses on thylakoid membrane proteins of WT, *cDEAP2-1*, *deap2-1* and *pam68-2* corresponding to the same amount of Chl, isolated from rosettes of low light grown plants. By using a WT dilution series, we could show that all PSII subunits were decreased ∼40% of WT in both *deap2-1* and *pam68-2* (Fig. 3A). When comparing both mutants with WT for other thylakoid complexes, on the basis of equal Chl, strong increases were found for the Cyt *b_6_f* and the NADPH-dehydrogenase (NDH) complexes as determined by PetA and NdhB antibodies, respectively and a mild increase for LHCII as detected by the Lhcb1 antibody. The chloroplast ATP synthase and PSI as detected by AtpB and PsaA antibodies, respectively, showed mild decreases. In parallel, PSII, PSI and Cyt*b_6_f* complexes were quantified spectroscopically (Fig. 3B). The results from this analysis supported the quantification by immunoblot analysis and a scheme summarizing the results from both methods is displayed in Fig 3C. In all experiments, the *cDEAP2-1* line showed comparable results to WT. The apparent increases in Cyt *b_6_f* in the mutants was due to data normalization on equal Chl basis. When normalized to leaf area, the content of Cyt *b_6_f* was unaltered in *deap2-1* and pam68-2 compared to WT, while PSII content was further reduced to 35% (Supplemental Table S3).

**Figure 3:**
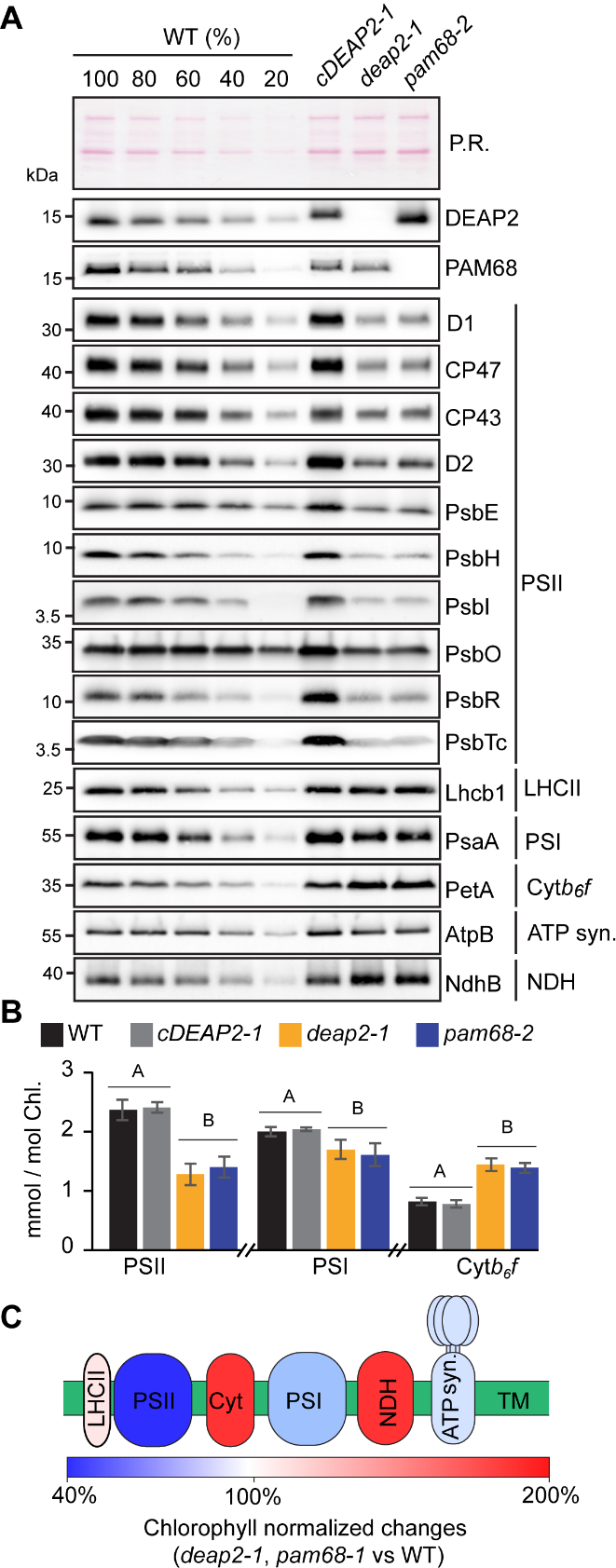
The *deap2* mutant contains similarly reduced PSII levels as *pam68*. **(A)** Immunoblot analysis of thylakoid protein corresponding to the same amount of chlorophyll (Chl) from WT, *cDEAP2-1*, *deap2-1* and *pam68-2* using specific antibodies against DEAP2 (1 µg Chl) and PAM68 (5 µg Chl), several subunits of photosystem II (PSII: D1, CP47, CP43, D2, PsbE, PsbH, PsbO, PsbR, PsbTc, all 1 µg Chl and PsbI, 5µg Chl) and representative subunits of light harvesting complex II (LHCII: Lhcb1, 1 µg Chl), photosystem I (PSI: PsaA, 1 µg Chl), cytochrome *b_6_f* complex (Cyt: PetA, 1 µg Chl), ATP synthase (AtpB, 1 µg Chl) and NADPH dehydrogenase complex (NDH: NdhB, 2 µg Chl). Ponceau red (P.R.) served as a control for equal loading. Numbers left of the immunoblots indicate molecular weight in kDa of protein standards. (**B)** PSII, PSI and Cyt*b_6_f* were quantified spectroscopically in isolated thylakoids. Averages are shown for n=5-7 ± SD. The capital letters above the graphs denote the two significantly different groups (p < 0.05) as determined by one-way ANOVA and Holm-Sidak multiple comparison test. **(C)** Scheme summarizing the relative Chl-normalized change in thylakoid membrane (TM) complex composition in both *deap2-1* and *pam68-2* compared with WT.

Together the quantification experiments performed on plants grown in low light conditions show that PSII is the thylakoid complex that is most drastically affected by loss of DEAP2 and that levels of all analyzed thylakoid membrane complexes are similarly affected in the *deap2-1* and *pam68-2* mutants.

### Ribosome profiling reveals that loss of DEAP2 or PAM68 have mild but similar effects on chloroplast transcript accumulation and translation

Both DEAP2 and PAM68 are integral thylakoid proteins that are required for the accumulation particularly of PSII. To further investigate the function of both DEAP2 and PAM68 in the biogenesis of plant PSII and distinguish between potential effects on transcript accumulation and translation, we performed chloroplast ribosome profiling (Fig. 4A (Trösch et al., 2018)). Changes in chloroplast RNA levels or ribosomal footprints (translation output) between mutants and WT were analyzed by a microarray-based method (see Material and Methods). All observed changes in transcript accumulation and translation output between the mutants and wild-type controls were below 2-fold indicating that neither DEAP2 nor PAM68 are likely to have a primary function in the expression of chloroplast-encoded PSII subunits or any other chloroplast gene (Fig. 3A). The analysis of differential changes compared with WT by volcano plots (Supplemental Fig. S3) revealed a number of significant, below 2-fold differences that were very similar between *deap2-1* and *pam68-2* (Fig 4B, Supplemental Tables S4-S7). On RNA level, both mutants showed significantly increased accumulation of multiple chloroplast transcripts encoding for subunits of PSII and the Cyt*b_6_f* complex, and decreased accumulation of *rbcL*. The calculation of translation efficiencies by normalizing translation output with RNA levels revealed a significant decrease for at least one subunit each of Cyt *b_6_f* (*petA*), PSI (*psaA*, *psaB*) and the ATP synthase (*atpB*) in both mutants. PSII transcript accumulation was significantly upregulated for *psbB* (CP47), *psbD* (D2)*, psbH* and *psbT*, while translation efficiency was decreased for *psbE* and *psbH* compared with WT.

**Figure 4:**
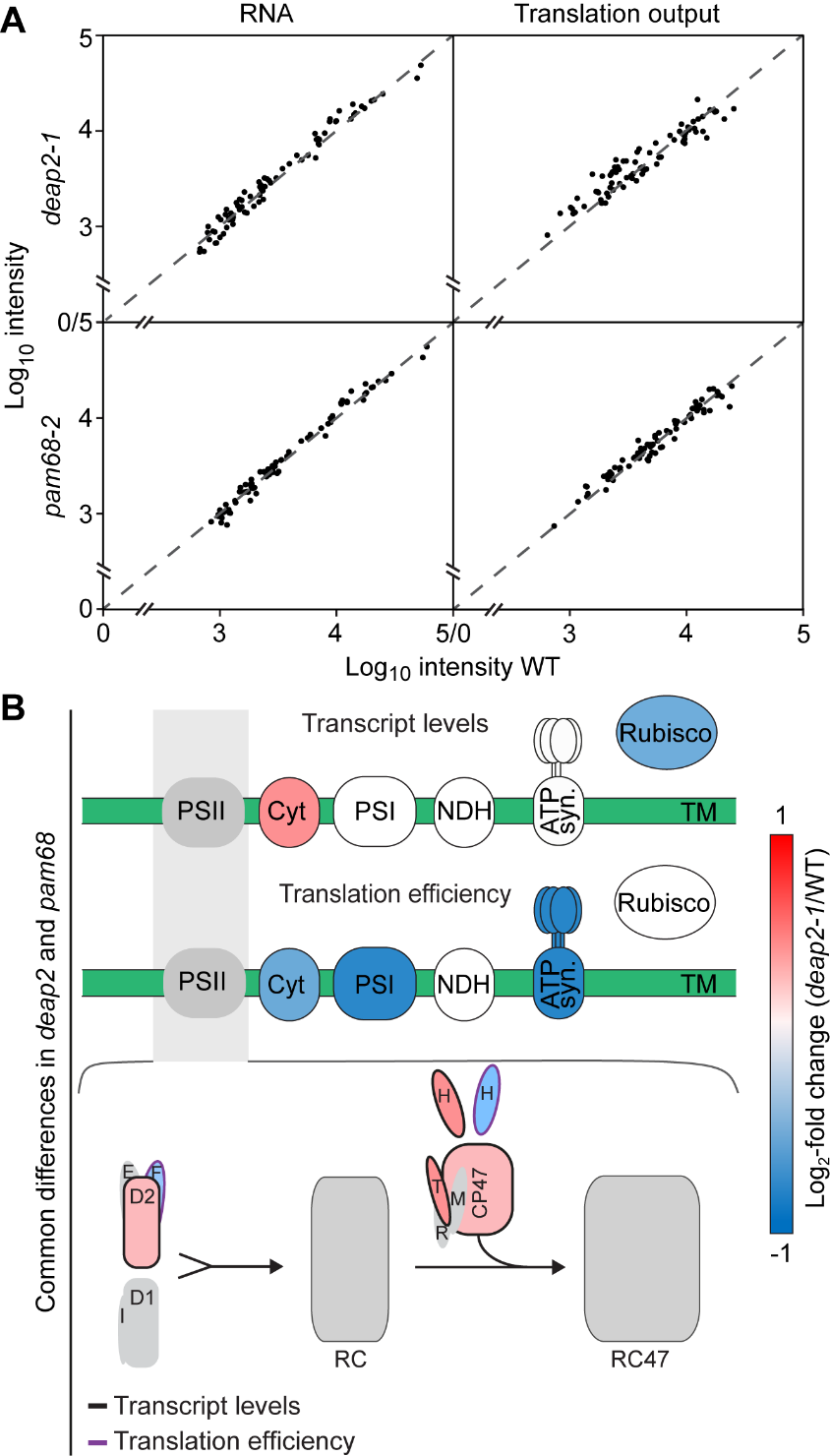
Levels of chloroplast-encoded transcripts and ribosomal footprints are slightly but similarly affected in *deap2* and *pam68* mutants. **(A)** RNA (left) and ribosomal footprints (right, translational output) of chloroplast transcripts from rosettes of *deap2-1* (top) and *pam68-2* (bottom) relative to WT. All values, without any specific outliers, cluster around the dotted line, which signifies equal abundance in the mutant vs. WT. **(B)** Graphical representation of common significant differences in both *deap2-1* and *pam68-2* mutants compared with WT for a single photosynthetic complex (upper panel, for values see Supplemental Table S7 and volcano plot in Supplemental Fig. S3). For coloring of complexes according to the color scale on the right, following genes were chosen: Transcription – Cyt *b_6_f* (*petB*) and Rubisco (rbcL); Translation efficiency – Cyt *b_6_f* (*petA*), PSI (*psaA*) and ATP synthase (*atpB*). The lower scheme displays the different steps of photosystem II assembly and significant differences in transcript levels (black outline) or translation efficiencies (purple outline) of assembled chloroplast-encoded subunits of photosystem II (PSII) between both *deap2-1* and *pam68-2* and WT. (Cyt, Cyt *b_6_f*; PSI, photosystem I; NDH, NADPH dehydrogenase complex; ATP syn., ATP synthase; Rubisco, ribulose-1,5-bisphosphate carboxylase/oxygenase; TM, thylakoid membrane).

Together, the data reveal similar effects of the *DEAP2* and *PAM68* lesions on chloroplast transcript accumulation and translation. These effects were rather small, demonstrating that both proteins are not essential for transcription, transcript processing or translation of chloroplast-encoded PSII subunits (or any other chloroplast gene). These comparably mild but common differences in transcript levels and translation efficiencies could be explained by regulatory feedback loops that are engaged to counteract the negative effects of both mutants on PSII accumulation. Here, the decrease in the translation efficiency of subunits encoding PSI and the ATP synthase may contribute to the lower accumulation of either complex in the thylakoid membrane of both mutants compared with WT (Fig. 3A-C).

### DEAP2 affects PSII assembly similarly to PAM68 and is likewise not associated with PSII assembly intermediates

To obtain insight into thylakoid complex assembly in the *deap2* mutant and compare it with *pam68*, we solubilized thylakoid membrane protein complexes with n-dodecyl-β-D-maltoside (DDM) and fractionated them by blue-native (BN)-PAGE (Fig. 5A).

**Figure 5:**
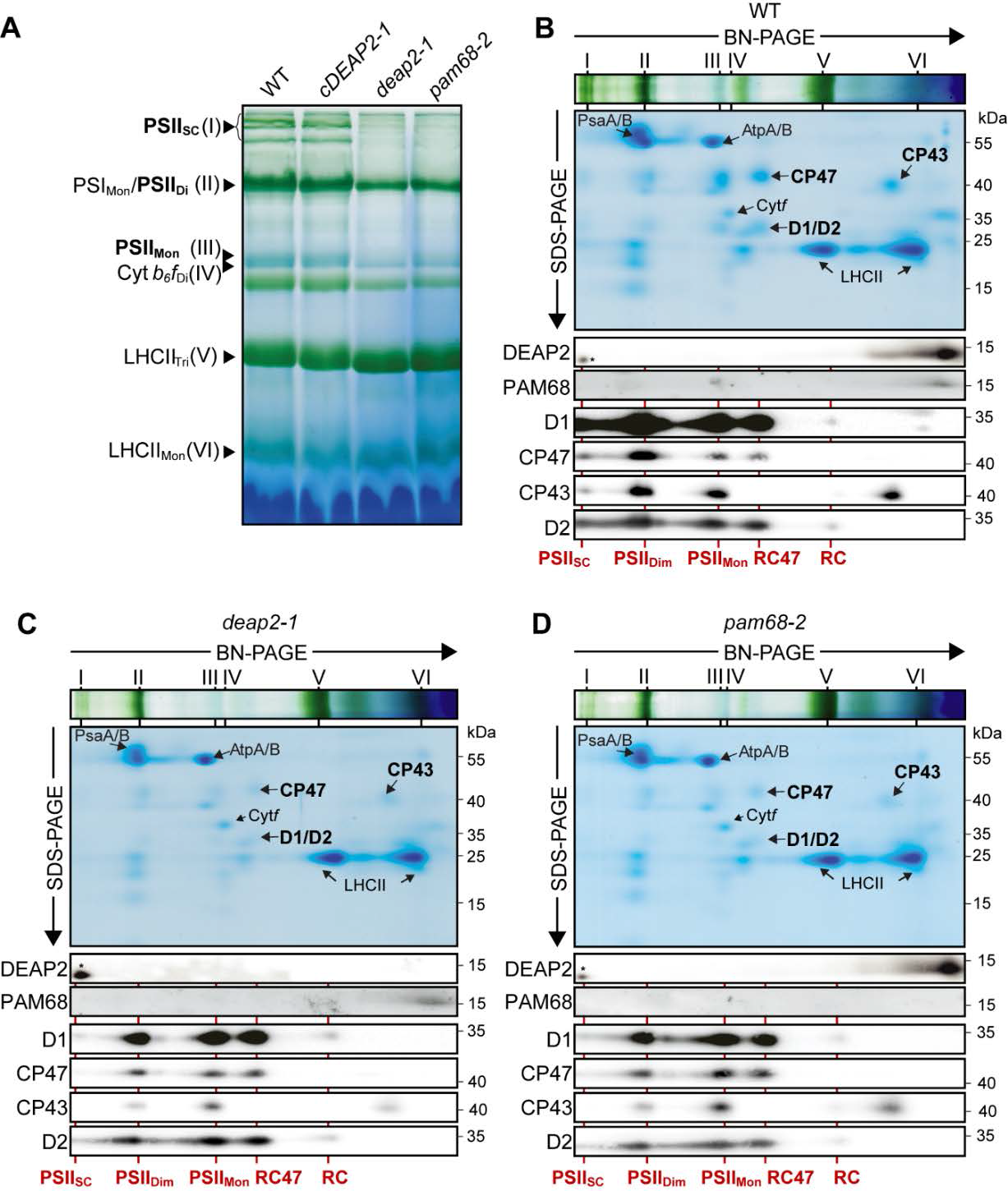
While the reaction center assembly intermediate accumulates to WT levels, higher molecular mass PSII assembly stages are reduced in *deap2* similar to *pam68*. **(A)** BN-PAGE analysis of thylakoid protein complexes. Thylakoid membranes of WT, *cDEAP2-1*, *deap2-1* and *pam68-2* were solubilized with 1 % (w/v) DDM at a concentration of 1 mg chlorophyll/ ml and fractionated by BN-PAGE. Visible bands were assigned according to previous reports (Armbruster et al., 2010) and results shown below: photosystem (PS) II supercomplexes (PSII_SC_; band I), PSI monomer and PSII dimer (PSI_Mon_/PSII_Di_; band II), PSII monomer (PSII_Mon_; band III), dimeric cytochrome *b_6_f* complex (Cyt *b_6_f*_Di_, band IV), trimeric light harvesting complex (LHC) II (LHCII_Tri_; band V) and monomeric LHCII (LHCII_Mon_; band VI). PSII complexes are highlighted in bold. (**B-D)** Thylakoid complexes on BN-PAGE slices of WT (B), *deap2-1* (C) and *pam68-2* (D) were separated into their subunits by SDS-PAGE. Gels were either stained with Coomassie or further processed for immuno-detection of DEAP2, PAM68 and the PSII core proteins D1, CP47, CP43 and D2 using specific antibodies. Roman numerals on top of the BN-slices indicate complexes as shown in (A). Red lines and labels indicate PSII assembly stages (RC, reaction center; RC47, RC+CP47). Numbers right of the immunoblots indicate molecular weight in kDa of protein standards.

Both WT and *cDEAP2-1* had a similar pattern of complex distributions with a strong blueish band at the expected size of monomeric PSII, running just above the Cyt*b_6_f* dimer, a strong green band at the size of the PSI monomer and PSII dimer and a number of green bands corresponding to the high molecular mass PSII supercomplexes (SC). These PSII containing bands were all markedly and similarly reduced in the *deap2-1* and *pam68-2* mutants, while LHCII trimers were slightly increased. Next, complexes were fractionated into their subunits by SDS-PAGE and visualized by Coomassie staining. This analysis supported the specific decrease of PSII, as only weak signals could be detected in the *deap2-1* and *pam68-1* mutants compared with WT and *cDEAP2-1* of the PSII core subunits D1, D2, CP43 and CP47 (Fig. 5C-D, Supplemental Fig. S4). Additionally, immunoblot analyses were carried out on the BN second dimensions using specific antibodies against DEAP2 and PAM68 as well the four PSII core subunits D1, D2, CP43 and CP47. Both DEAP2 and PAM68 were mainly found to migrate as monomers or small molecular weight complexes and were not present in PSII complexes or its assembly intermediates. We found both D1 and D2 to accumulate in WT and the mutant lines to similar levels in an early low abundant complex, which we assigned to the RC assembly intermediate. In accordance with the Coomassie stained gels, levels of D1, D2, CP47 and CP43 were lower in both *deap2-1* and *pam68-2* compared with WT and *cDEAP2-1*. In both mutants, most signal of the PSII subunits was found in two assembly stages, one lacking CP43, which we assigned to the RC47 assembly intermediate and the other one representing the PSII monomer. For WT and *cDEAP2-1* the strongest signal was found in the PSII dimer and some also in PSII SC. Both knock-out mutants showed very little signal of PSII subunits in the PSII SC.

The 2D BN-PAGE analysis revealed that DEAP2 like PAM68 does not co-migrate with PSII assembly intermediates, but is present in the thylakoid membrane as a monomer or in small molecular weight complexes. Both DEAP2 and PAM68 are required for the full accumulation of PSII assembly intermediates downstream of the RC, particularly for the final functional complexes, the PSII dimer and SC. Together, the results strongly support that DEAP2 has a function in PSII assembly, potentially at the same step as PAM68, which is the assembly of RC47 from RC.

### Both DEAP2 and PAM68 are required for the formation of RC47 from RC

To monitor the dynamics of PSII assembly, we labelled newly synthesized chloroplast encoded proteins radioactively. For this, discs of young leaves of WT, *deap2-1* and *pam68-2* (see Supplemental Fig. S5 for more details) were first infiltrated with cycloheximide to block cytosolic translation, then with 35S-labelled methionine and floated on infiltration medium at low light intensity for 20 min. From half of the leaf discs, thylakoids were isolated (pulse), while the second half was washed and infiltrated with non-labelled methionine and incubated in low light for another 20 min before isolation of thylakoid membranes (chase). Thylakoid protein complexes were solubilized using DDM, separated by 2D-BN-PAGE and the radioactive signal of labelled proteins was detected (Fig. 6A).

**Figure 6:**
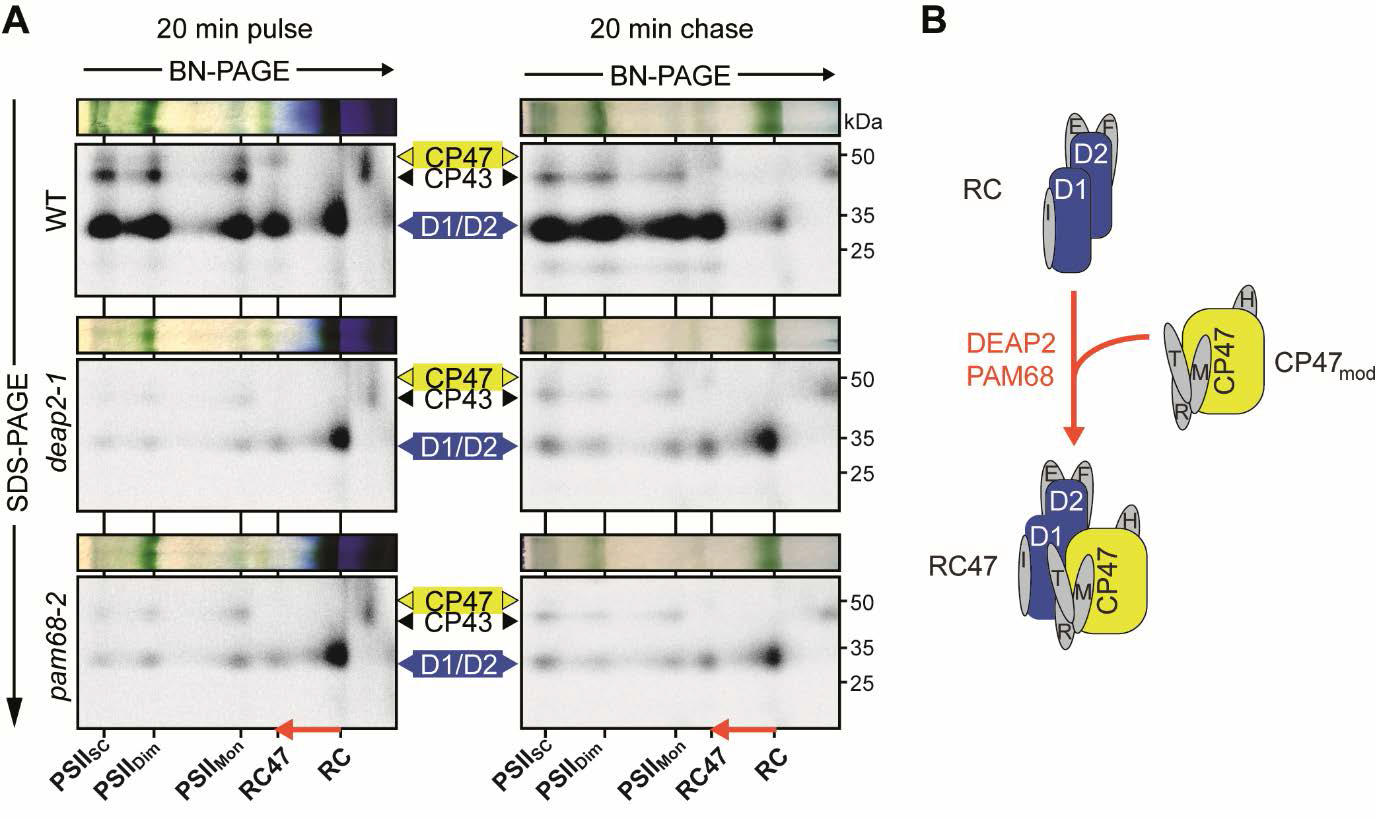
PSII assembly in *deap2* stalls at the RC stage exactly as in *pam68*. **(A)** Analysis of the dynamics of photosystem II (PSII) assembly by labeling newly synthesized chloroplast proteins with [^35^S] methionine and separating thylakoid complexes and denatured subunits by 2-dimensional gel electrophoresis. Leaf discs from WT, *deap2-1* and *pam68-2* were pulse labeled for 20 min in the presence of cycloheximide and thylakoids were directly processed, or leaf discs were first infiltrated again with non-radioactive methionine to be able to chase [^35^S] methionine incorporation into higher PSII assembly stages (supercomplexes (PSII_SC_), dimers (PSII_Di_), monomers (PSII_Mon_), reaction center (RC) and RC+CP47 (RC47). The red arrow shows the transition from the RC to the RC47 assembly intermediates. Numbers right of the auto-radiographs indicate molecular weight in kDa of protein standards. **(B)** Scheme illustrating the transition from RC to RC47, of which efficient processing is dependent on the presence of both DEAP2 and PAM68.

In WT, mainly D1/D2, CP43 and CP47 were labelled with similar amounts in all respective PSII assembly stages after the pulse. After the non-radioactive chase, the signal of the RC intermediate was nearly gone and more label was found in the PSII assembly stages of higher molecular mass. In both, the *deap2-1* and the *pam68-1* mutants, we found that the RC D1/D2 signal was comparable to WT after the pulse. Only very few label was observable in further higher molecular PSII assembly stages in these mutants. In contrast to WT, the non-radioactive chase did not greatly deplete the level of label in the RC and signal in the higher molecular weight complexes remained low.

Together, these analyses reveal that *deap2-1* and *pam68-2* show identical defects in the dynamics of PSII assembly. Loss of either protein dramatically impairs the process downstream of the RC formation (Fig. 6B).

### Plants that lack both DEAP2 and PAM68 cannot make functional PSII

Because loss of DEAP2 has a comparable effect on PSII assembly as loss of PAM68, we investigated the possibility that both proteins act cooperatively in this process. For this, we generated double mutants of *deap2-1* and *pam68-2*. No double mutants were identified from F2 plants grown on soil, suggesting that these plants could not grow photo-autotrophically. Thus, the selection was continued on sucrose-containing agar plates and double mutants could be isolated. These plants were white, small and had no measurable Fv/Fm (Fig. 7A), indicating that plants lost the capacity to grow photo-autotrophically, because they lacked PSII. To confirm this, immunoblot analyses were performed on total protein extracted from leaf discs of WT, single mutants and the *deap2-1 pam68-2* double mutant (Fig. 7B). While the content of the Actin control was comparable between the genotypes, levels of PSII subunits were below detection level in the *deap2-1 pam68-2* double mutant. Analyzed subunits for the Cyt*b_6_f* complex, PSI and ATP synthase were lower as compared to single *deap2-1* and *pam68-2* mutants, but were detectable at around 50% of WT. Thus, other thylakoid complexes accumulated to much higher levels than PSII in the double mutant, supporting that PSII assembly is the main defect in the double mutant. To control for the specificity of this identified genetic interaction between *deap2* and *pam68*, we additionally crossed *pam68-2* with *lpa1-2*, a mutant with impaired integration of D1 in the thylakoid membrane. These double mutants were displayed an intermediate Fv/Fm value between the slightly lower *pam68-2* and higher *lpa1-2* (Supplemental Fig. S6 A, B). The loss of both PAM68 and LPA1 in these plants was confirmed by immunoblotting using specific antibodies against both proteins (Supplemental Fig. S6 C).

**Figure 7:**
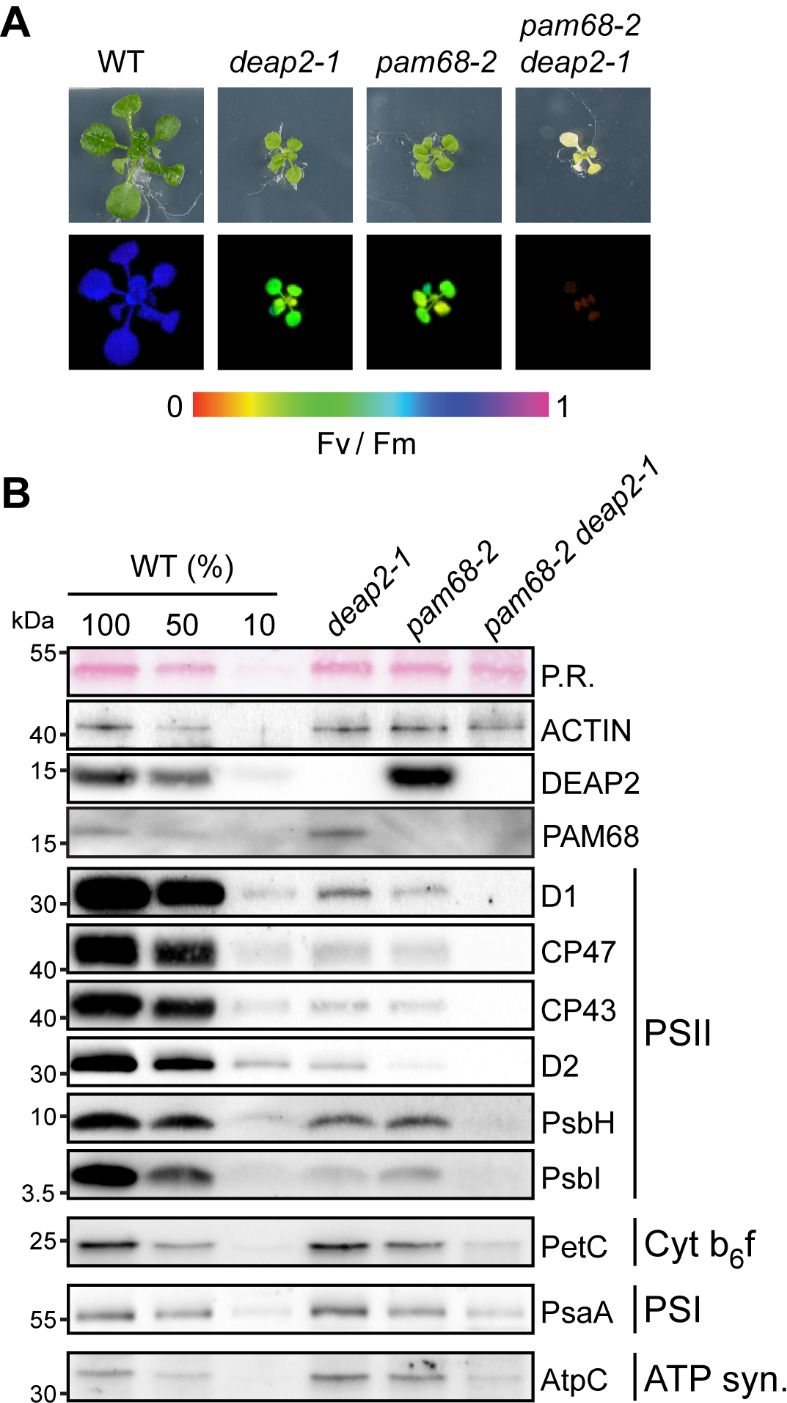
Plants lacking both DEAP2 and PAM68 do not accumulate any functional PSI. **(A)** Plants lacking both DEAP2 and PAM68 (cross of *pam68-2* and *deap2-1*) are only viable on MS medium supplemented with 2 % sucrose. Pictures (upper panel) and false color maximum photosystem (PS) II quantum yield (Fv/Fm) images (lower panel) of 4-week-old WT, *deap2-1*, *pam68-2* and *pam68-2 deap2-1* double mutants grown at 50 µmol photons m^−2^ s^−1^ and 12 h/ 12h dark light cycles. Signal intensities for Fv/Fm are given by the false color scale below the panels. **(B)** Immunoblot analysis of total protein extracts from plants as in (A). Samples were fractionated by SDS-PAGE and blots were probed with specific antibodies against DEAP2, PAM68, several subunits of PSII (D1, CP47, CP43, D2, PsbH, PsbI), Cyt b6f (PetC), PSI (PsaA) and the ATP synthase (AtpC). Ponceau red (P.R.) stained RbcL on the membrane before detection and Actin were used to control for equal protein loading. Numbers left of the immunoblots indicate molecular weight in kDa of protein standards.

Together, the data demonstrate that simultaneous absence of DEAP2 and PAM68 leads to a complete disruption of PSII assembly. Possible explanations for this could be a cooperative or a redundant function of the two proteins during the assembly of RC47 from RC and the CP47 module.

### DEAP2 overexpression can partially rescue the loss of PAM68, but not vice versa

To further test for a potential redundancy of DEAP2 and PAM68 in the PSII assembly process, we generated mutant lines, in which the other protein was overexpressed as a Myc-tagged version. For control, we additionally generated *PAM68* overexpression plants in the *pam68-2* background. From five positive transformants each, as detected by the Myc antibody, we selected three with varying levels of overexpression and performed further experiments on the two with the lowest and highest expression (Fig. 8A, Supplemental Fig. S7).

**Figure 8:**
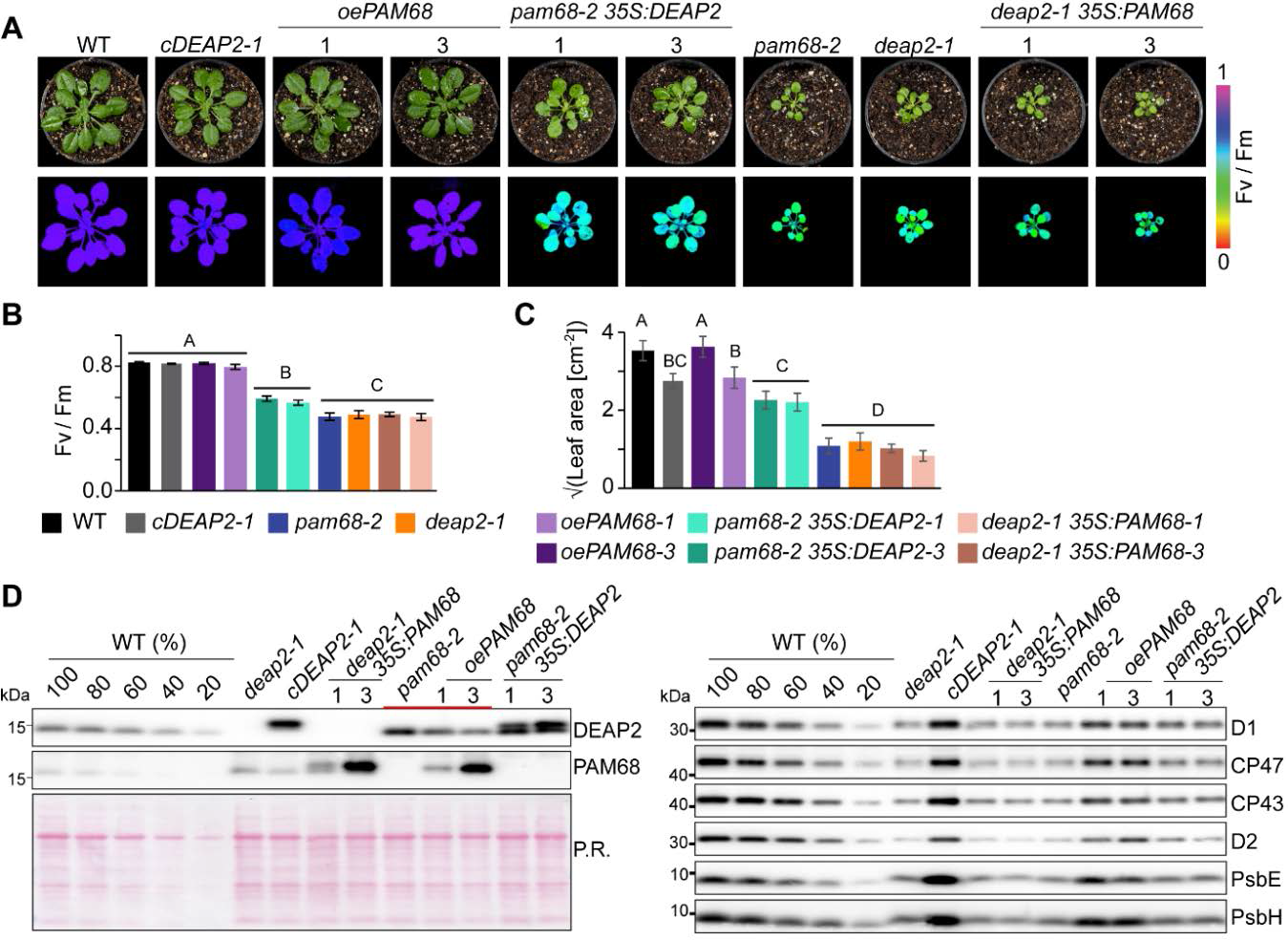
DEAP2 can partially compensate for loss of PAM68 but not vice vers. **(A)** Pictures (upper panel) and false color images of the maximum quantum yield of photosystem (PS)II (Fv/Fm, lower panel) of six-week-old WT, *deap2-1*, *cDEAP2-1*, *pam68-2*, *deap2-1* overexpressing *PAM68* (*deap2-1 35S:PAM68*), *pam68-2* overexpressing *PAM68* (*oePAM68*) or *DEAP2* (*pam68-2 35S:DEAP2*). All overexpressed proteins carry a c-terminal MYC tag. Three independent lines with varying amounts of protein expression were selected (Supplemental Fig. 7). Shown here are lines #1 and #3 with lowest and highest Myc-tagged protein levels as determined by immunoblotting. Signal intensities for Fv/Fm are given by the false color scale below the panels. **(B-C)** Fv/Fm (B) and leaf area (C, shown as square root) from n=6-10 genotypes as shown in (A) with error bars indicating standard deviation. Capital letters above graphs indicate groups with significant differences between genotypes (p < 0.05) as determined by one-way ANOVA and Holm-Sidak multiple comparison test. **(D)** Immunoblot analysis of total protein extracts using specific antibodies against DEAP2 and PAM68 (left panel) and PSII subunits (right panel). The accumulation of subunits of other thylakoid complexes is shown in Supplemental Fig. 7. Ponceau red (P.R.) staining of one membrane prior to immunodetection is shown as a loading control.

Overexpression of *PAM68* in *pam68-2* complemented the low growth and Fv/Fm phenotype, but did not have any effects in *deap2-1*, while overexpression of *DEAP2* in the *pam68-2* rescued these two phenotypes to some degree (Fig. 8B,C). These *pam68-2* plants with 2-fold levels of DEAP2 had significantly higher Fv/Fm and larger leaf area compared with *pam68-2* alone. To investigate whether this was due to higher levels of PSII accumulating in the *pam68-2 35S:DEAP2* lines, we performed immunoblot analyses on multiple subunits and saw generally a slight increase compared with *pam68-2* (Fig. 8D). By comparing the signals of DEAP2 and PAM68 in the different lines, we observed that DEAP2 decreased with increasing PAM68 levels.

To obtain information about the relationship between DEAP2 and PAM68 on protein level, we quantified their relative abundances (Fig. 9A). For equal amounts of Myc-tagged protein, we loaded 1x *cDEAP2-1,* which had similar DEAP2 levels as WT (Supplemental Fig. S8), and 4x *oePAM68-3*, which had five-fold higher PAM68 levels than WT (Supplemental Fig. S7, 8). This result shows that DEAP2 is ∼20x more abundant than the PAM68.

**Figure 9:**
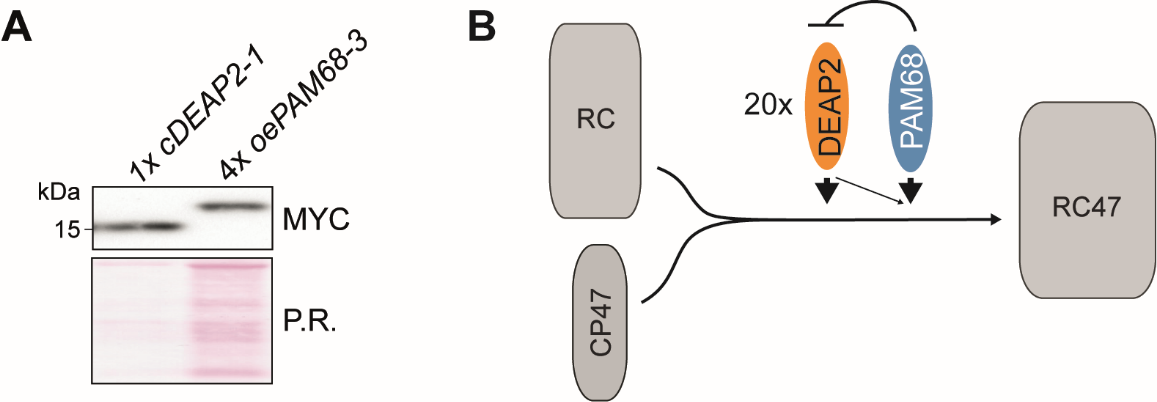
DEAP2 has 20-fold higher protein levels than PAM68 and model of function. (A) Quantification of relative PAM68 and DEAP2 levels. Total protein extracts of 1x *cDEAP2-1* (WT-levels) and 4x*oePAM68-3* (five-fold overexpression) yield the same Myc signal. (C) Schematic model of the transition from reaction center (RC) PSII assembly stage to the RC+CP47 (RC47) intermediate. This assembly step is facilitated by DEAP2 and PAM68, whereby DEAP2 is 20-fold more abundant than PAM68. Increasing PAM68 levels lead to decreasing DEAP2 accumulation and DEAP2 to can partially compensate for lack of PAM68

## Discussion

### DEAP2 is a eukaryote-specific factor that cooperates with PAM68 in the progression of the RC to the RC47PSII assembly intermediate

This study has uncovered DEAP2 as a new PSII assembly factor, which together with the known assembly factor PAM68 facilitates the formation of the RC47 intermediate from RC and the CP47 module. We reveal that lack of DEAP2 and PAM68 have nearly identical effects on all stages of thylakoid biogenesis and PSII assembly. Both, the *deap2* and *pam68* mutants show comparable defects in the accumulation of thylakoid complexes with a strong reduction in PSII subunits and assembled PSII dimers as well as super-complexes (Fig. 3, 5). Decreases in the levels of PSI and the ATP synthase in the mutants compared with WT can be explained by the downregulation of some of their subunits at the translational level, potentially involving a negative feedback loop in response to low PSII accumulation (Fig. 4). Consistent with this, it has previously been shown that many PSII biogenesis mutants with pronounced reductions in their PSII content display parallel, but milder decrease in PSI levels. For example, in tobacco mutants with reduced *psbD* mRNA abundance, a decrease in PSII content to 20% of wild-type levels resulted in a reduction of PSI content to 35% of wild type levels (Fu et al., 2020). Also, Arabidopsis mutants specifically affected in PSII assembly show a concomitant reduction in PSI accumulation (Meurer et al., 1998; Peng et al., 2006; Link et al., 2012; Li et al., 2019). Together, this further supports that the decreased PSI content observed here is a consequence of altered PSII abundance, and not attributable to any additional function of DEAP2 or PAM68 related to PSI.

PSII assembly of both mutants is impaired at the RC assembly intermediate. Lack of either protein strongly decreases the progression from RC to RC47. The PAM68 orthologue in cyanobacteria was previously shown to interact with ribosomes and proposed to promote CP47 synthesis and co-translational insertion of Chl molecules (Bucinska et al., 2018). Such a function of PAM68 could explain why the assembly process is slowed down at the RC also in plants. However, the incorporation of radioactive label into newly synthesized thylakoid CP47 protein was not affected in plants lacking PAM68 (Armbruster et al., 2010). The ribosome profiling analyses of this report again argue against an effect of Arabidopsis PAM68 on the translation of CP47, as we could not observe any differences in *psbB* translation efficiencies between WT and *pam68* (Fig. 4, Supplemental Fig. S3, Supplemental Table S. 7). We did find however a decrease in *psbH* translation efficiencies in both mutants, which may indicate a function of these proteins related to the assembly of PsbH together with CP47. For cyanobacteria it was previously suggested that PsbH shares the same CP47 binding site with PAM68 and replaces PAM68 after the insertion of Chl into CP47 (Bucinska et al., 2018).

Together, our results suggest that the assembly of RC47 from the RC intermediate is not as highly conserved between plants and cyanobacteria as previously thought. Our data do not support a similar function of plant PAM68 in CP47 synthesis, maturation and assembly as has been reported for cyanobacteria. This may indicate that the PAM68 protein has adopted a different function throughout the evolution from cyanobacteria to plants. Its general role in RC47 formation however has been maintained. Differently to PAM68, DEAP2 is not conserved in cyanobacterial genomes, but is only present in the genomes of the green lineage of photosynthetic eukaryotes and thus represents a eukaryotic addition to the PSII RC47 formation process (Karpowicz et al., 2011). For the plant PSI assembly process, it has previously been described that in addition to highly conserved factors, it also involves proteins that represent eukaryotic inventions (Albus et al., 2010; Schöttler et al., 2011).

### DEAP2 and PAM68 appear to have complementary functions

We demonstrate that DEAP2 can at least partially compensate for the loss of PAM68, but not vice versa. The two-fold amplification of DEAP2 levels in the *pam68* background by expression of a second *DEAP2* copy from the 35S promoter, partially rescued the low Fv/Fm and growth phenotypes of the *pam68* mutant (Fig. 8B). However, particularly PSII subunit accumulation and Fv/Fm as a read-out for functional PSII were still strongly reduced compared with WT, demonstrating that even when DEAP2 is present in excess, PAM68 is still required for an efficient PSII assembly process. Given that DEAP2 is 20 times more abundant than PAM68 (Fig. 9), this points to a function of PAM68 which cannot be entirely replaced with DEAP2. Our results rather suggest that an excess of DEAP2 buffers against the negative consequences that loss of PAM68 has on PSII assembly. The absence of functional PSII assembly in the double mutant is also consistent with this hypothesis. A four-fold excess of PAM68 cannot ameliorate the negative effects of DEAP2 loss on PSII functionality. This may be due to overexpressed PAM68 being present in sub-stoichiometric amounts compared with DEAP2 and thus not accumulating to high enough levels for entirely taking over the function of DEAP2. In this case, however, we would expect at least a partial rescue of the *deap2* phenotype by PAM68 overexpression, which is not the case. The inability of PAM68 overaccumulation to compensate for the loss of DEAP2 rather supports the hypothesis that both proteins have distinct functions. Interestingly, levels of DEAP2 appeared to decrease with increasing PAM68 levels (Fig. 8D). A 4-fold increase in PAM68 caused a slight decrease in DEAP2 levels up to 80% of WT. In line with the 20-times higher levels of DEAP2 in the thylakoid membrane compared with PAM68, this may suggest a stoichiometric destabilizing effect of PAM68 on DEAP2. The instability of a larger pool of DEAP2 is supported by the observation that *deap2-1* plants overexpressing DEAP2 from the 35S promoter can only accumulate WT levels of DEAP2. Also, in *pam68-2* mutants, DEAP2 levels only reach a maximum of two-fold compared with WT.

## Conclusion

While multiple PSII assembly factors have been reported, exact mechanistic insight into how most of these factors promote PSII biogenesis is still lacking. The presented study breaks grounds for further investigations into how (i) RC47 assembly differs between cyanobacteria and plants and (ii) DEAP2 and PAM68 contribute to this process mechanistically. In particular, molecular understanding of how DEAP2 stabilizes the transition from RC to RC47 even in the absence of PAM68 may reveal strategies to enhance and accelerate the safe assembly of plant PSII, but potentially also other multi-partite complexes with highly reactive co-factors.

## Material and Methods

### Plant Material and Growth Conditions

The Salk T-DNA insertion lines *deap2-1* (SALK_048033), *pam68-2* (SALK_044323, (Armbruster et al., 2010)), *lpa1* (SALK_124398), herein referred to as *lpa1-2*, were obtained from NASC (Alonso et al., 2003) and genotyped using specific primers (Supplemental Table 6). The insertion in *deap2-1* was confirmed by sequencing the flanking region of the insertion. To generate plant expression constructs, the *DEAP2* or *PAM68* coding regions were amplified from cDNA using gene-specific primers adding a sequence encoding for a C-terminal MYC-tag and inserted into the binary vector pEarleyGate100 (pEG100, (Earley et al., 2006)) linearized with *Xba*l and *Xho*l restriction enzymes downstream of the *35S* promoter by using Gibson Assembly (Gibson et al., 2009). Additionally, the *DEAP2* cDNA and the sequence encoding for YFP were introduced into pEG100 in the same way to obtain a protein fusion of DEAP2 to YFP at its C-terminus. Homozygous *deap2-1* and *pam68-2* plants were transformed by floral-dip as described previously (Clough and Bent, 1998). Transgenic plants (*cDEAP2*, *oePAM68*, *deap2 35S:PAM68*, *pam68 35S:DEAP2*) were selected based on their resistance to BASTA and expression of the fusion protein was detected by immunoblot analysis using an antibody against the Myc tag.

Arabidopsis plants were grown in pots on soil (Stender, Schermbeck, Germany) containing 1 g/l Osmocote or on Murashige and Skoog medium with 0.8 % agar supplemented with 2 % sucrose (w/v) at ∼50 µmol photons m^−2^ s^−1^, 20 °C/ 16 °C, 60%/ 75% relative humidity in 12h/ 12h day/ night cycles. Experiments were performed on 6 to 8-week-old WT, *cDEAP2*, *oePAM68* and 10 to 12-week-old *deap2-1*, *pam68-2*, *lpa1*, *deap2 35S:PAM68*, *pam68 35S:DEAP2*.

### Transient expression of DEAP2-YFP fusions in *Nicotiana benthamiana* leaves

The *Agrobacterium tumefaciens* strain GV3101 transformed with the DEAP2−YFP constructs was resuspended in induction medium (10 mM MgCl_2_, 10 mM MES-KOH pH 5.6, 150 μM acetosyringone) to an OD600 of 0.5. After 2 h at 28 °C, suspensions were inoculated onto sections of well-watered *N. benthamiana* (tobacco) leaves by injecting into the bottom side (Blatt and Grefen, 2014). Transfected plants were grown for 2 d before leaf discs were analyzed for the subcellular localization of the YFP fluorescence signal. For microscopy, the Leica TCS SP5 instrument was used with 63x/1.4 objective and water immersion. Fluorophores were excited by using an argon laser at 514 nm, YFP fluorescence was detected between 524 and 582 nm, and Chl fluorescence between 618 and 800 nm.

### *In-silico* analyses

Sequence alignment and phylogenetic reconstructions were done using ClustalW (https://www.genome.jp/tools-bin/clustalw). Sequences were retrieved from Phytozome v13 (https://phytozome-next.jgi.doe.gov). Alignment and phylogenetic reconstructions were performed using the function “build” of ETE3 3.1.2 as implemented on the GenomeNET (https://www.genome.jp/tools-bin/ete). The tree was constructed using FastTree v2.1.8 with default parameters. The Arabidopsis DEAP2 chloroplast transit peptide (cTP) and transmembrane domains was predicted using TargetP (Armenteros et al., 2019) and DeepTMHMM (Hallgren et al., 2022), respectively. The protein structure of DEAP2 (without cTP) was predicted by Alphafold (Jumper et al., 2021) or I-TASSER (Zhou et al., 2022), and overlayed using PyMols *super* function (Delano, 2002). The I-TASSER model quality was evaluated by estimated RMSD, estimated TM-score, structural alignments against known protein structures from PDB and the I-TASSER own quality parameter C-score. Additionally, the online tool MolProbity (Davis et al., 2007) was used for stereo-chemical quality (Ramachandran) analysis of all predicted models.

### Protein isolation, gel electrophoresis and immunodetection

Total protein was extracted from pulverized leaf material using sample buffer (0.1 M Tris pH 6.8, 24 % glycerol (v/v), 8 % SDS, 5 % mercapto-ethanol (v/v), 0.02 % bromophenol blue (w/v)) and heating at 40 °C for 30 min. Thylakoid membranes were isolated as described previously (Armbruster et al., 2014), and total Chl content was estimated according to Porra et al., 1989 (Porra et al., 1989). For SDS polyacrylamide gel electrophoresis (PAGE), total protein extracts or thylakoid samples incubated with sample buffer were resolved on a 16% polyacrylamide gel containing 4M urea (Schagger, 2006). For blue native-PAGE (BN-PAGE) thylakoid membranes at a concentration equal to 1 mg of total Chl/ mL were solubilized with 1 % β-n-dodecyl-d-maltoside (DDM), and separated on a 6-12 % gradient gel according to (Peng et al., 2008). The BN-PAGE was further fractionated by SDS-PAGE. For this, BN-PAGE gel strips were first incubated with 66 mM Na_2_CO_3_, 2 % SDS and 50 mM DTT at 50°C for 30 min to denature proteins. After electrophoresis, gels were either stained with Coomassie blue (R-250, 40% methanol, 7% acetic acid, 0.05% Coomassie blue R-250) or used for immunoblotting analyses. For immunoblot analyses with specific antibodies, gels were blotted onto nitrocellulose and visualized with Ponceau Red (0.1 % Ponceau S (w/v) in 5 % (v/v) acetic acid) prior to immunodetection. Membranes were blocked with 50 mM Tris, 10 mM NaCl, 0.05 % (v/v) Tween 20, 0.2 % (v/v) Triton™ X-100 and 5% (w/v) nonfat dry milk and incubated with antibodies against D1 (AS10704), CP47 (AS04038), CP43 (AS111787), D2 (AS06146), PsbE (AS06112), PsbH (AS06157), PsbI (AS06158), PsbO (AS06142-33), PsbR (AS05059), Lhcb1 (AS01004), Lhcb2 (AS01003), PsaA (AS06172), PetA (AS06119), PetC (AS08330), AtpB (AS05085), AtpC (AS08312), RbcL (AS03037), NdhB (AS164064), Tic40 (AS10709), Actin (AS214615-10), Myc (AS153034), all obtained from Agrisera (Sweden), as well as PAM68 (PHY2289A) and PsbTc (PHY0350A) obtained from PhytoAB and diluted according to manufacturer’s instructions. For immunodetection of DEAP2, an antibody was produced by Pineda Antikörper-Service (Germany) targeting the C-terminal end of the protein (peptide sequence: VEAANKEAQEQDKRDGFL) and diluted 1:1000. For secondary antibodies coupled to the horse-radish peroxidase, a mouse- or rabbit-specific antibody was used from Agrisera (against Actin; AS09627) or Sigma-Aldrich (A8275), respectively, at a dilution of 1:10,000. Signals were detected using the SuperSignal™ West Femto Chemiluminescence-substrate (ThermoFisher) and the G-Box-ChemiXT4 (Syngene).

### Chloroplast isolation, fractionation and chaotropic salt treatment of isolated thylakoids

Intact Arabidopsis chloroplasts were isolated via a two-step Percoll gradient centrifugation (Kunst, 1998). In brief, the homogenized leaf tissue was fractionated on the gradient and intact chloroplasts were collected from the interphase. Chloroplasts were fractionated into envelope, thylakoids and stroma via a three-step sucrose gradient centrifugation after being hypotonically lysed in 10mM Tris/HCl, pH 8.0 and 1mM EDTA, pH 8.0, at a Chl concentration of 2 mg/ mL (Li et al., 1991). For salt washes of thylakoids, isolated thylakoids were resuspended in 50 mM HEPES/KOH, pH 7.5, at a Chl concentration of 0.5 mg/ mL (Karnauchov et al., 1997). Extraction with 2 M NaCl, 0.1 M Na_2_CO_3_, 2 M NaSCN, or 0.1 M NaOH was performed for 30 min on ice, soluble and membrane proteins were separated by centrifugation for 10 min at 10,000*g* and 4°C, and immunoblot analysis was performed on both fractions.

### Spectroscopic plant analysis and photosynthetic complex quantification

The MAXI IMAGING-PAM (Walz, Effeltrich, Germany) was used for Chl *a* fluorescence analysis. Minimal fluorescence F0, and maximal fluorescence Fm of 30 min dark-acclimated plants were measured before and during the application of a 0.8 s saturating light pulse, respectively. The maximum yield of PSII Fv/Fm was calculated as (Fm-F0)/Fm. PSI measurements were performed with the plastocyanin-P700 version of the Dual-PAM instrument (Schöttler et al., 2007), which allows the deconvolution of absorbance changes arising from plastocyanin and PSI. Plants were directly taken from the controlled environment chambers and measured without dark adaptation. The fraction of PSI reaction centers limited at the donor side, Y(ND), was determined according to (Schreiber and Klughammer, 2016). All measurements were performed in the middle of the light period.

Thylakoid membranes were isolated as previously described (Schöttler et al., 2004). Their Chl content and a/b ratio were measured in 80% (v/v) acetone according to Porra et al. (1989). The contents of PSII and the Cyt *b_6_f* were determined from difference absorbance signals of Cyt *b_559_* (PSII) and the cytochromes *b_6_* and *f*, respectively. To improve the optical properties of the samples, thylakoids equivalent to 50 µg Chl mL^−1^ were de-stacked in low-salt medium supplemented with 0.03% (w/v) DDM (Kirchhoff et al., 2002). All cytochromes were fully oxidized by the addition of 1 mM potassium hexacyanoferrate (III). Then, 10 mM sodium ascorbate was added to reduce the high-potential form of cyt b_559_ and cytochrome f. Finally, the addition of 10 mM sodium dithionite reduced the low potential form of cyt b_559_ and the two b-type hemes of cytochrome b_6_. Using a V-750 spectrophotometer equipped with a custom-made head-on photomultiplier (Jasco GmbH, Pfungstadt, Germany), at each of the three redox conditions, 10 absorbance spectra were measured between 575 and 540 nm with a spectral bandwidth of 1 nm and a scanning speed of 100 nm min^−1^, and subsequently averaged. Difference spectra were calculated by subtracting the spectrum measured in the presence of hexacyanoferrate from the ascorbate spectrum, and by subtracting the ascorbate spectrum from the spectrum measured in the presence of dithionite, respectively. Finally, a baseline calculated between 540 and 575 nm was subtracted from the signals. Then, the difference spectra were deconvoluted as previously described (Kirchhoff et al., 2002). PSI was quantified from light-induced difference absorbance changes of P_700_. Thylakoids equivalent to 50 µg Chl mL^−1^ were solubilized in the presence of 0.2% (w/v) DDM. After the addition of 10 mM sodium ascorbate as electron donor and 100 µM methylviologen as electron acceptor, P_700_ was fully photo-oxidized by applying a light pulse (250 ms length, 2000 µmol photons m^−2^ s^−1^ intensity). Measurements were performed with the Pc-P700 version of the Dual-PAM instrument (Heinz Walz GmbH). This instrument was also used to measure light response curves of PSI parameters on intact leaves. The donor-side limitation Y(ND) of PSI was calculated according to Klughammer and Schreiber (2016) (Schreiber and Klughammer, 2016). Finally, based on the Chl content of the investigated leaves, all complex contents were re-normalized to a leaf area basis.

### Immunoprecipitation of DEAP2 and following MS analysis

Thylakoids were solubilized in 50 mM HEPES pH 8.0; 330 mM Sorbitol; 150 mM NaCl; 4 mM PMSF containing 1% β-DM for 10 min on ice. The supernatant was recovered after centrifugation and incubated with magnetic beads at 4 °C overnight. Beads that were made specific for DEAP2 by attaching the antibody to Protein G-coupled beads (Dynabeads, ThermoFisher) according to protocol, or Myc-antibody coupled beads (Myc-trap, ChromoTek) were used. After incubation, the beads were washed five times with the same buffer containing 0.2 % β-DM, before samples were further processed. For MS analysis, proteins were reduced with TCEP, alkylated with chloroacetamide, and digested with trypsin for 14 h at 37°C. The peptide mixture was purified and desalted on C_18_ SEP-Pak columns (Waters). Measurements were performed on a Q Exactive HF or Exploris 480 mass spectrometer coupled with a nLC1200 nano-HPLC (Thermo Scientific). Quantitative analysis of MS/MS measurements was performed with the MaxQuant software (Cox and Mann, 2008) and the Andromeda search engine was used to annotate peptide sequences using the Arabidopsis TAIR10 genome annotation database.

### Ribosome profiling

Plant material was harvested 1 h after the start of the photoperiod from six-week (WT) and 10-week (mutants) old plants (at comparable developmental stage), immediately frozen in liquid nitrogen and stored at −80 °C until further processing. For each line, three replicates were used, consisting of 10 independent plants. The ribosome profiling was done as previously described (Zoschke and Barkan, 2015; Trösch et al., 2018).

Ribosome footprints and total RNA were prepared, labelled and hybridized as described previously (Trösch et al., 2018; Schuster et al., 2020; Ghandour et al., 2023). Briefly, 4 μg ribosome footprint RNA and 3.5 μg total RNA from *deap2-1*, *pam68-2* and wild-type plants were chemically labelled with Cy3 and Cy5, respectively, and hybridized onto identical custom microarrays as described (Trösch et al., 2018). These microarrays contain 50-nt probes that cover chloroplast protein-coding regions and UTRs in high density (one 50mer probe starting every 30 nt within these regions; each probe is represented in four technical replicate spots on each microarray). Transcript and ribosome profiling data were processed as described previously (Schuster et al., 2020; Gao et al., 2022). In brief, background-subtracted probe signals lower than 100 were set to zero. Probes that showed signal saturation were excluded from further analysis. Signal intensities of all probes covering protein-coding regions (in all experiments and replicates) were normalized to the average signal intensities of probes covering protein-coding regions in the three wild-type datasets (Supplemental Tables S4,5). For each chloroplast reading frame the average signal intensity values based on the signal intensities of all probes located in a given reading frame was calculated to determine the fold-change differences in comparison to wild-type data. Significances of changes in gene expression were calculated by Student’s *t*-test and the derived *p* values were adjusted for multiple testing as described previously (Benjamini and Hochberg, 1995).

### *In-vivo* labeling of newly synthesized chloroplast proteins

Labeling of thylakoid protein was performed as previously described (Armbruster et al., 2010) on young leaf discs from six-week (WT) and 10-week (mutants) old plants (at comparable developmental stage) at 3 h after the start of the photoperiod (Supplemental Fig. S5). Briefly, leaf discs with a 0.5 cm diameter were vacuum-infiltrated with 20 µg/ ml cycloheximide in 10 ml of 10 mM Tris, 5 mM MgCl2, 20 mM KCl, pH 6.8, and 0.1 % (v/v) Tween 20 and incubated for 30 min to block cytosolic translation. For pulse labeling, the leaves were infiltrated again with the same solution containing 1 mCi of [^35^S] methionine and transferred into ambient light (∼10 µmol photons m^−2^ s^−1^) for 20 min. Then, thylakoid isolation and BN-PAGE were performed as described above. Following the pulse labeling, leaves were incubated again with 10 mM of unlabeled methionine for 20 min in light (chase labeling) and processes further as described above. Signals were detected using Storage phosphor screens (Fujifilm) and visualized using the Amersham Typhoon RGB Biomolecular Imager (GE Healthcare Life Sciences).

## Data availability

MS raw data were deposited via JPOST (Moriya et al., 2019). The data is available under the following link and will be made publicly available upon publication. URL: https://repository.jpostdb.org/preview/208626461964527a25c90d0. For access key contact the corresponding author.

## Accessions

DEAP2 sequences from Arabidopsis (At3g51510), *Glycine max* (Glyma.03G158200), *Oryza sativa* (Os05g41190), *Sorghum bicolor* (Sobic.009G179800), *Zea mays* (ZMPHB47.06G250900), *Sphagnum maghellanicum* (Sphmag03G126700) and *Physcomitrium patens* (Pp3ca2_25230), *Volvox carteri* (Vocar.0018s0093) and *Chlamydomonas reinhardtii* (Cre16.g679300) and PAM68 and LPA1 from Arabidopsis (At4g19100 and At1g02910, respectively).

## Funding and Acknowledgement

JMK, UA, MAS, RZ and RB received funding from the Max Planck society and RZ and IF from the Deutsche Forschungsgemeinschaft (DFG # 437345987 and 445970965, 469950637, respectively). The authors thank Paulina Heinkow for excellent technical assistance within the MS Proteomics Unit Biology of Plants of the University of Münster.

## Author contributions

MJF selected T-DNA insertion lines, made complementation lines and tested the antibody. OSI generated additional mutants by CRISPR-Cas employing expertise from RB. SK generated double mutants. JMK performed most experiments with the help of LJ and NW. JE and IF performed MS analyses and interpretations. MAS and WT quantified complexes and pigments and measured photosynthetic parameters. IG and RZ performed ribosome profiling. TR predicted protein structure. JMK, UA, RZ, MAS and DS interpreted data. UA wrote manuscript together with JMK and help from all authors.

