## Supplemental Fig for "A eukaryote-specific factor mediates an early step in the assembly of plant photosystem II"

Keller et al.,

Supplemental Figures S1- S8

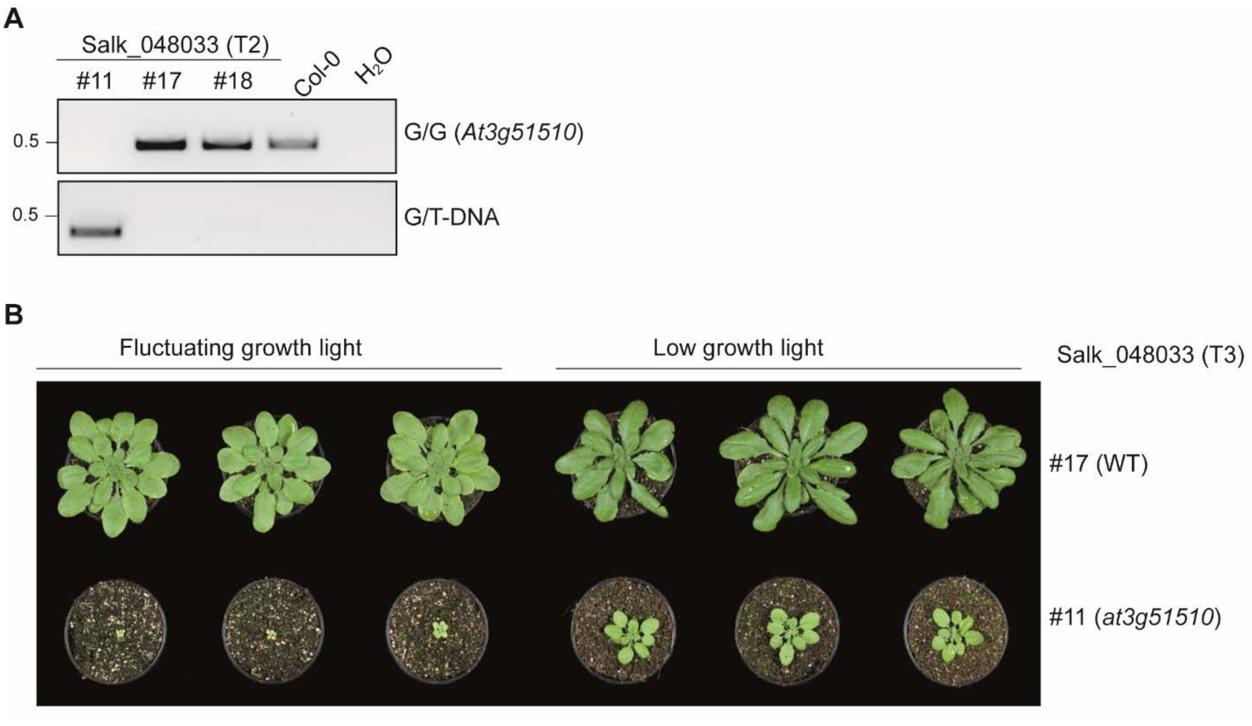


**Supplemental Figure S1: Selection of *At3g51510* T-DNA insertion lines**
**(A)** T2 plants from the Salk_048033 seed stock were genotyped for the T-DNA insertion using primers specific for either the gene (G) alone or together with a left border primer of the T-DNA, and one homozygous mutant and two wild type (WT) plants were identified and grown for the production of T3 seeds. **(B)** Seed from T3 # 11 (*at3g51510* mutant) and #17 (WT) were sown out under fluctuating (144 repetitions of 1 min 90 µmol photons $m^{-2}$ $s^{-1}$, 4 min 900 µmol photons $m^{-2}$ $s^{-1}$) and low growth light (90 µmol photons $m^{-2}$ $s^{-1}$, both conditions with 12 h/ 12 h light/ dark cycles) for five weeks before pictures were taken.

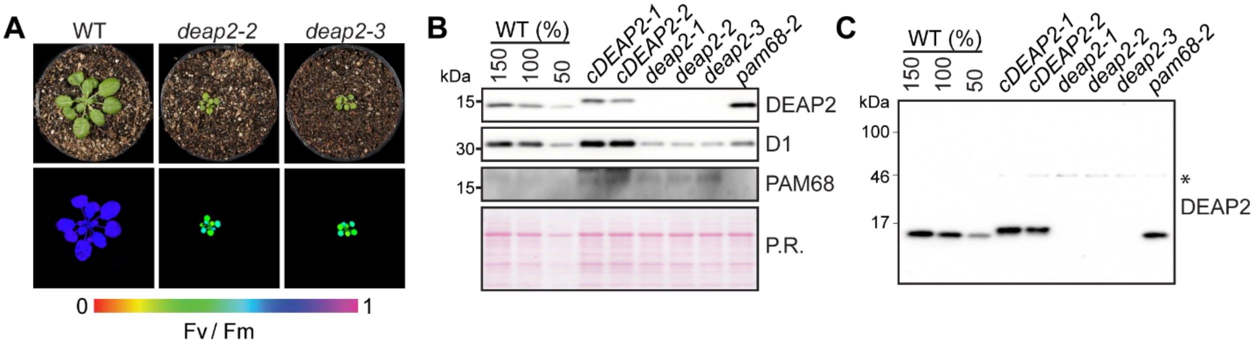


**Supplemental Figure S2: CRISPR-Cas deletion lines of *DEAP2***

**(A)** Pictures (top panel) and PSII maximum quantum yield (Fv/Fm) as false color images taken with the Imaging-PAM (bottom panel) of six-week-old Col-0 (wild type, WT), *deap2-2* and *deap2-3*, generated by using the CRISPR-Cas system, grown at 50 µmol photons $m^{-2}$ $s^{-1}$ with a 12/12 h light/dark cycle. Signal intensities for Fv/Fm are given by the false color scale below the panels. (**B)** A specific DEAP2 antibody was generated against the 15 most C-terminal amino acids of the protein. Immunoblot analysis of total protein extracts of WT, the complemented lines *cDEAP2-1*, *cDEAP2-2*, the three *deap2* mutants *deap2-1*, *deap2-2*, *deap2-3* and *pam68-2* reveals that the antibody specifically recognizes DEAP2 with an upwards shifted signal in the Myc tagged *cDEAP2* lines and a missing signal in all three mutants. An additional immunoblot analysis shows a reduction of the PSII-D1 subunit below 0.5 of WT protein levels in all three *deap2* mutants as well as the *pam68* mutant. Detection of PAM68 demonstrates its accumulation in the *deap2* mutant and absence in the *pam68* mutant. Ponceau staining (P.R.) of the membrane prior to immunodetection is shown as loading control. Numbers left of the immunoblots indicate molecular weight in kDa of protein standards. (**C)** Immunodetection of DEAP2 over the whole membrane as in (B). Besides the specific DEAP2 signal below the 17 kDa marker band, a weak unspecific signal is visible at about 45 kDa (indicated by an asterisk).

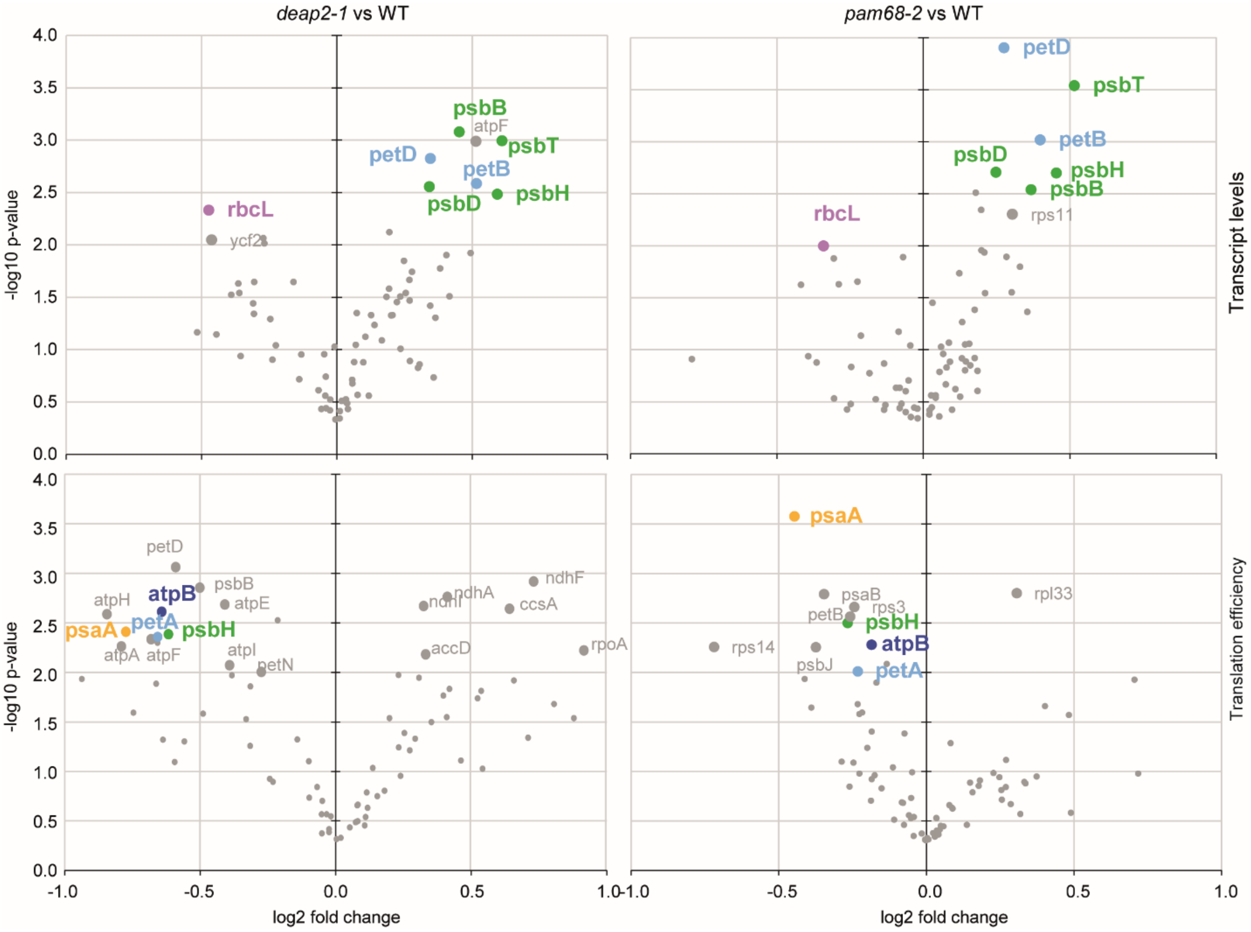


**Supplemental Figure S3: Vulcano plot analysis of changes in transcript and translation efficiency between *deap2-1* and *pam68-2* compared with WT**

The data from the ribosome profiling experiments were analyzed by volcano plot to identify common and distinct changes in *deap2-1* and *pam68-2* compared with WT. Those data points are labelled for which the -log10 p-value was above 2 (corresponding to p-value < 0.01) and log-fold change was <-0.3 and >0.3 for transcript levels and <-0.19 and >0.19 for translational efficiency. Such data points found for a specific transcript in both the *deap2-1* and *pam68-2* analysis are shown in bold and in color, with light blue for genes encoding for the Cyt $b_6f$ complex, purple for Rubisco, green for photosystem II (PSII), dark blue for the ATP synthase and orange for PSI.

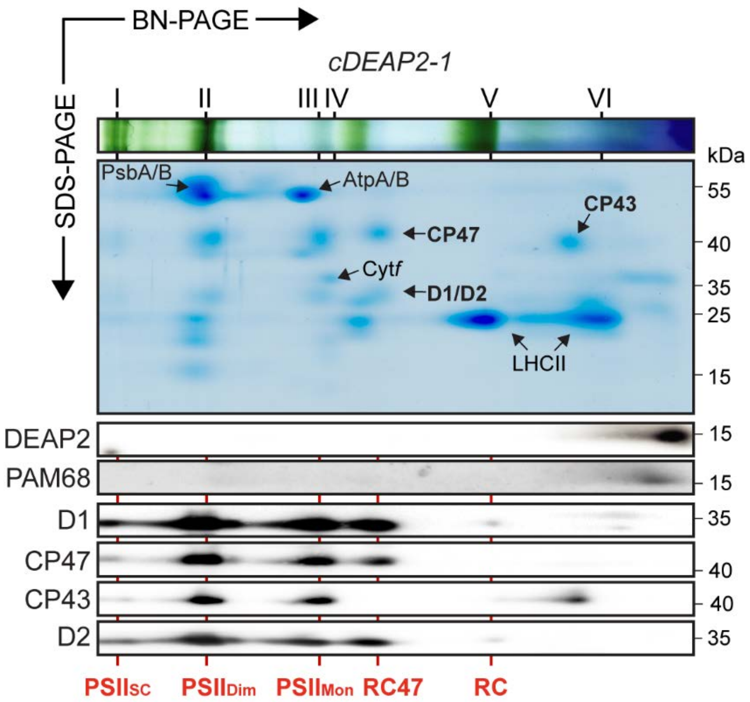


**Supplemental Figure S4: 2D-BN-PAGE analysis of solubilized *cDEAP2-1* thylakoid complexes**

Thylakoids were isolated and solubilized as described for the other lines in Fig. 5. Solubilized thylakoid complexes were fractionated by blue native (BN)-PAGE. Roman numerals on top indicate complexes according to previous reports (Armbruster et al., 2010). Photosystem (PS) II supercomplexes (band I), PSI monomer and PSII dimer (band II), PSII monomer (band III), dimeric cytochrome $b_6f$ complex (band IV), trimeric light harvesting complex (LHC) II (V) and monomeric LHCII (band VI). Coomassie staining of the second dimension is shown below the BN-PAGE, indicating the signals of PSII core proteins D1/D2, CP47, CP43, as well as PsaA/B, AtpA/B, Cyt*f* and LHCII. Numbers right of the immunoblots indicate molecular weight in kDa of protein standards. DEAP2, PAM68 and the PSII core proteins D1, CP47, CP43 and D2 were detected by immunoblot analysis and signals are shown below the Coomassie stained gel. Red lines and labels indicate PSII assembly stages (RC, reaction center; RC47, RC+CP47; $PSII_{Mon}$, PSII monomers; $PSII_{Dim}$, PSII dimers; $PSII_{SC}$, supercomplexes).

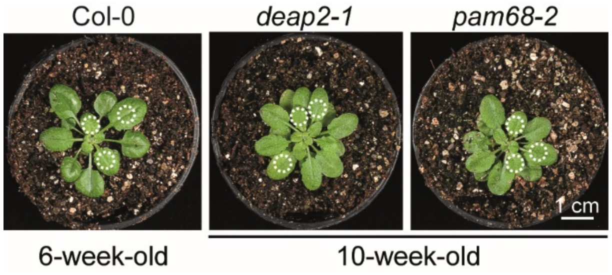


**Supplemental Figure S5: Leaf discs used for *in-vivo* labelling**

Discs for *in vivo* labelling were taken from plants of comparable leaf size, 6-week-old wild type (WT) and 10-week-old *deap2-1* and *pam68-2*. Dashed lines around circles indicate the size of the discs (5 mm diameter)

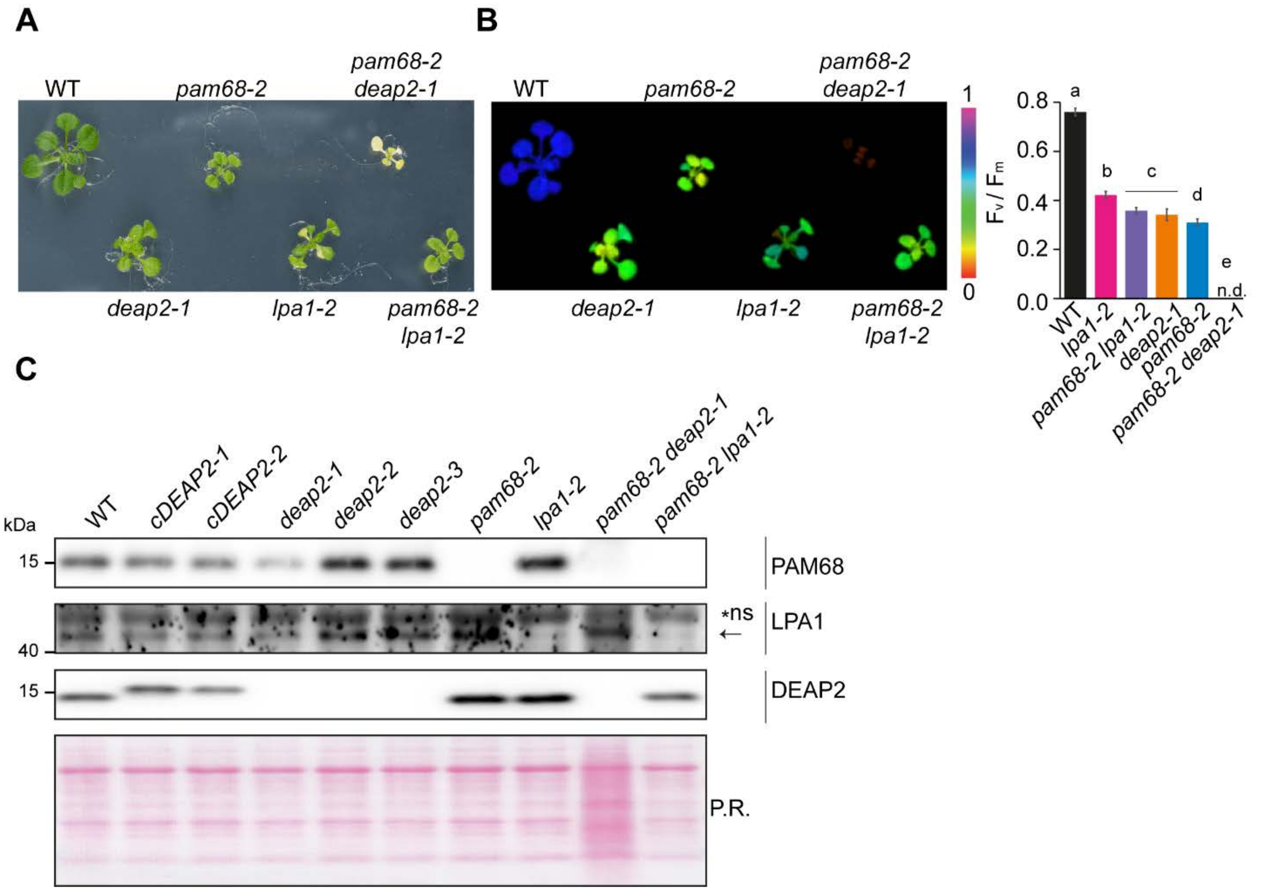


**Supplemental Figure S6: Double mutant of *pam68* with *deap2* but not with *lpa1* show a lack of functional PSII**

**(A-B)** Plants lacking both PAM68 and DEAP2 (cross of *pam68-2* and *deap2-1*) are only viable on MS medium supplemented with 2 % sucrose and completely lack functional PSII. Pictures (A) and false color images of maximum photosystem (PS) II quantum yield (Fv/Fm, B) are shown for 4-week-old WT, *deap2-1*, *pam68-2* and *pam68-2 deap2-1* double mutants grown at 50 µmol photons $m^{-2}$ $s^{-1}$ and 12 h/ 12h dark light cycles. Signal intensities for Fv/Fm are given by the false color scale. On the right is a bar graph of Fv/Fm determined from n=5 ±SD. Different small letters above graphs indicate significant differences between raw data of the different genotypes ($p < 0.05$) as determined by one-way ANOVA and Holm-Sidak multiple comparison test. n.d.=not detectable **(C)** Immunoblot analysis on total protein extract isolated from plants grown on MS medium supplemented with 2 % sucrose using specific antibodies against PAM68, LPA1 and DEAP2. Ponceau staining (P.R.) of the membrane prior to immunodetection is shown. Numbers left of the immunoblots indicate molecular weight in kDa of protein standards.

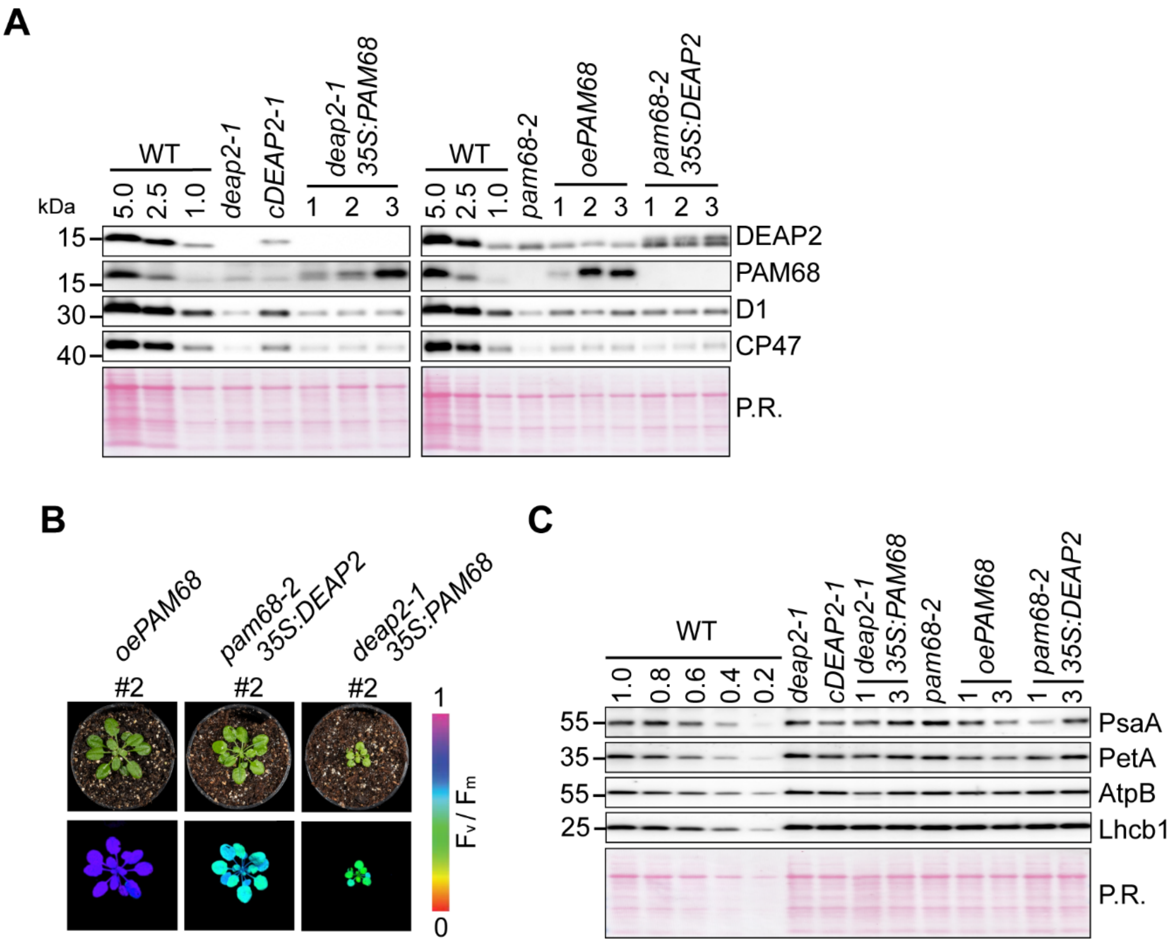


**Supplemental Figure S7: Selection and characterization of *DEAP2* and *PAM68* expressing mutants**

**(A)** Immunoblot analysis of total protein extracts from WT, *deap2-1*, cDEAP2-1 and three *deap2-1* lines overexpressing *PAM68* (left panel) and WT, *pam68-2* and three *pam68-2* lines each either overexpressing *PAM68* or *DEAP2* (right panel) using specific antibodies against DEAP2 and PAM68 as well as the PSII core subunits D1 and CP47. Ponceau staining (P.R.) of the membrane prior to immunodetection is shown. Numbers left of the immunoblots indicate molecular weight in kDa of protein standards. **(B)** Pictures (upper panel) and false color images of the maximum quantum yield of photosystem (PS)II (Fv/Fm, lower panel) of six-week-old *pam68-2* overexpressing *PAM68* (*oePAM68*) or *DEAP2* (*pam68-2 35S:DEAP2*) and *deap2-1* overexpressing *PAM68* (*deap2-1 35S:PAM68*) with intermediate protein levels (#2). All overexpressed proteins carry a c-terminal MYC tag. Signal intensities for $F_v/F_m$ are given by the false color scale below the panels. **(C)** Immunoblot analysis of total protein extracts from plants as shown in Fig.7 using specific antibodies against core subunits of PSI (PsaA), the cytochrome *b₆f* (PetA, Cyt *f*), the chloroplast ATP synthase (AtpB) and the light harvesting complex II (Lhcb1). Ponceau staining (P.R.) as shown in (A).

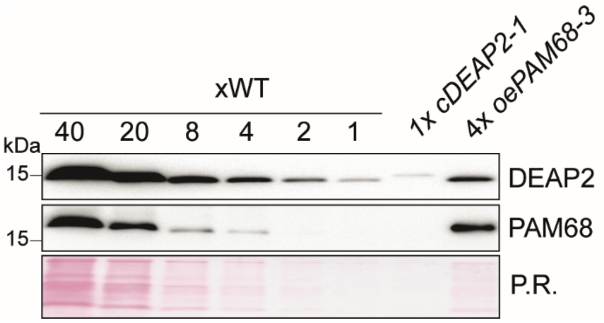


**Supplemental Figure S8: Relative quantification of DEAP2 and PAM68**

A WT dilution series of total protein extract was loaded together with *cDEAP2-1* and *PAM68-3* corresponding to the same Myc signal (Fig. 9) and DEAP2 and PAM68 were detected by immunoblot analysis.
